# Cisplatin-Induced Plasticity Drives Cross-Resistance to CDK4/6 Inhibitors and Reveals Targetable Vulnerabilities Through HDAC Inhibition in Esophageal Squamous Cell Carcinoma

**DOI:** 10.64898/2026.07.17.738886

**Authors:** Marta Ávalos-Moreno, Aurora Vigato, Fabiana Moresi, Diego Japón-Ruiz, Benjamin Beck, Xavier Bisteau

## Abstract

Esophageal squamous cell carcinoma (ESCC) accounts for 90% of esophageal cancer cases worldwide and carries a poor prognosis, with overall survival below 20%. Chemotherapy remains the standard of care, yet acquired resistance is a critical barrier and the programs that establish and stabilize it are still poorly defined. Using an *in vitro* model of cisplatin resistance, we show that sustained platinum exposure does not drive a single, uniform resistant state but a continuum of adaptive transcriptional states. Most resistant clones presented a drug-tolerant persister (DTP)-like phenotype with stemness traits and slow-cycling behaviour. The DTP phenotype persisted after cisplatin withdrawal and was accompanied by markers of resistance, suggesting consolidation into a stable, acquired-resistance state. Unexpectedly, the effect of this reprogramming extended beyond cisplatin. Resistant clones displayed impaired cytostatic control and failed to suppress cell cycle programs upon CDK4/6 inhibition (CDK4/6i), revealing collateral cross-resistance. Multiomic profiling revealed an attenuation of the immune and inflammatory signaling expected from CDK4/6i, upregulation of immune-evasive markers and a dysregulated IFN pathway activation at baseline in cisplatin-resistant clones, reflecting an immune-evasive state consequent to cisplatin-induced reprogramming. A signature-based approach identified HDAC inhibitors as candidates for reversing resistance, and functional validation confirmed that HDACi combined with cisplatin fully restored parental-level sensitivity while also enhancing CDK4/6i efficacy, supporting epigenetic reprogramming as a strategy to counteract cross-resistance. Together, our findings define cisplatin resistance in ESCC as a reversible, epigenetically encoded and immune-evasive state, and provide a mechanistic rationale for HDACi-based combinations to restore sensitivity to both cisplatin and CDK4/6 inhibitors.

**Statement of Significance:** Sustained cisplatin exposure drives esophageal squamous cell carcinoma toward a drug-tolerant persister-like, immune-evasive state that survives drug withdrawal and confers cross-resistance to CDK4/6 inhibition. HDAC inhibition reverses this state, restoring sensitivity to both agents and synergizing in chemotherapy-naive cells, supporting epigenetic intervention early in treatment.

## Introduction

Esophageal cancer remains among the deadliest cancers, being the 7th most common cause of cancer death and the 11th most common cancer^1^. While EC comprises two subtypes of cancer, esophageal squamous cell carcinoma (ESCC) accounts for 90% of EC cases worldwide^2^ and has been associated with a poor overall survival below 20% globally^3,4^. The current standard of care often includes platinum salts (i.e. carboplatin, cisplatin) in combination with radiotherapy or other chemotherapeutic agents. However, patients often develop resistance and relapse. It is therefore essential to gain a better understanding of the molecular landscape of ESCC to develop alternative therapeutic strategies.

Genomic analyses of ESCC have pictured a broad and heterogeneous landscape of oncogenic alterations^5,6^. Among these alterations, the cell cycle has appeared to be frequently deregulated, especially at the level of the cyclin D-CDK4/6-pRB axis, making this pathway a promising target for therapy^6,7^. Several CDK4/6 inhibitors (*palbociclib*, *ribociclib* and *abemaciclib*) have been FDA approved and are regularly used as first-line treatment for advanced ER+/HER2- breast cancer in combination with endocrine therapy^7^.

Our laboratory had previously demonstrated that CDK4/6 inhibition in naïve ESCC cell lines led to a similar response to that seen in breast cancer tumors, characterized by a strong cell cycle arrest and a pro-inflammatory, immunomodulatory response^8^. Nonetheless, recent clinical trials (NCT01037790, NCT03339843) testing the benefit of CDK4/6 inhibition as a monotherapy against advanced ESCC patients reported limited clinical activity. Interestingly, all enrolled were refractory to previous genotoxic chemotherapy (e.g. cisplatin). In this context, we wonder how genotoxic treatment may affect the tumoral molecular landscape and modulate their response to subsequent therapy with CDK4/6 inhibition in ESCC, an aspect that has not yet been addressed. Notably, many exploratory clinical trials initiated following results from naive models are performed in a second- or third-line setting, without elucidating the changes imposed by the first line of treatment in patients. It is therefore crucial to understand how genotoxic drugs like cisplatin affect tumor cells and their response to other therapies like CDK4/6 inhibition. Understanding these mechanisms may reveal strategies to rescue the sensitivity to CDK4/6i and lead to novel combination therapies.

In this study, we show that cisplatin exposure produces not a binary resistance switch, but a dynamic landscape of adaptive states, shaped predominantly by pre-existing cellular lineage. Most resistant clones converged on a DTP-like phenotype that remained stable after drug withdrawal, identifying DTP-like cells as a clinically relevant reservoir for relapse and refractory disease in ESCC patients completing platinum-based chemotherapy. This acquired resistance also reshaped response to subsequent therapies. Cisplatin-induced plasticity drove cross-resistance to CDK4/6 inhibitors (CDK4/6i) through a dual mechanism: loss of the cytostatic effect and attenuation of the interferon-driven inflammatory response associated to CDK4/6i efficacy. Using a signature-based drug-repurposing strategy, we identified HDACi as a state-reversal agent capable of re-sensitizing resistant cells to cisplatin and partially restoring CDK4/6i sensitivity. HDACi also synergized in chemotherapy-naive cells, positioning them as resistance-impairing agents when incorporated early into treatment. Collectively, this study reframes acquired cisplatin resistance as a dynamic, plastic process with direct clinical applications, and provides preclinical evidence supporting the rational design of a combination strategy with cisplatin, HDACi and CDK4/6i.

## Results

### Cisplatin exposure drives a continuum of adaptive resistant states rather than a single resistance mechanism in ESCC cells

While cancer cell lines may harbour intrinsic mechanisms of drug resistance, the adaptive resistance programs arising from sustained chemotherapy exposure remain incompletely understood. To address this, we established an isogenic *in vitro* model of cisplatin resistance in ESCC and studied the alterations following drug resistance. In order to establish cisplatin-resistant cell lines both pulsatile and continuous exposure paradigms were implemented, designed to reflect clinically relevant pharmacokinetics (**Fig. 1A**)^9–12^, as well as to recapitulate a broader spectrum of adaptative mechanisms arising from the same molecular background and to find converging mechanisms that could constitute major contributors to acquired resistance. Additionally, four parental cell lines differing in oncogenic alterations and baseline cisplatin sensitivity were selected (KYSE140, KYSE180, KYSE410 and TE6), so that the panel would span the genetic diversity of ESCC tumours alongside the adaptive variability sampled within each background. Cisplatin treatment consistently led to a significant increase in IC50 across all ESCC cell lines regardless of exposure method, with the largest fold-changes observed in the clones derived from the most sensitive parental lines, KYSE140 and KYSE180 (**Fig. 1B, C**). Despite differences in exposure modality (pulsatile or continuous), resistant clones derived from the same parental line generally exhibited highly overlapping transcriptional reprogramming profiles, with KYSE140RC2/RP2 and KYSE180RC2/RP2 sharing 75.1% and 80.8% of differentially expressed genes, respectively (**Fig. 2A; Sup. Fig. 1A-C**). Notably, transcriptional divergence was more pronounced across different cellular backgrounds (**Fig. 2A**). This data suggested that alterations following cisplatin exposure may be more influenced by underlying biology rather than treatment regimen.

**Figure 1.**
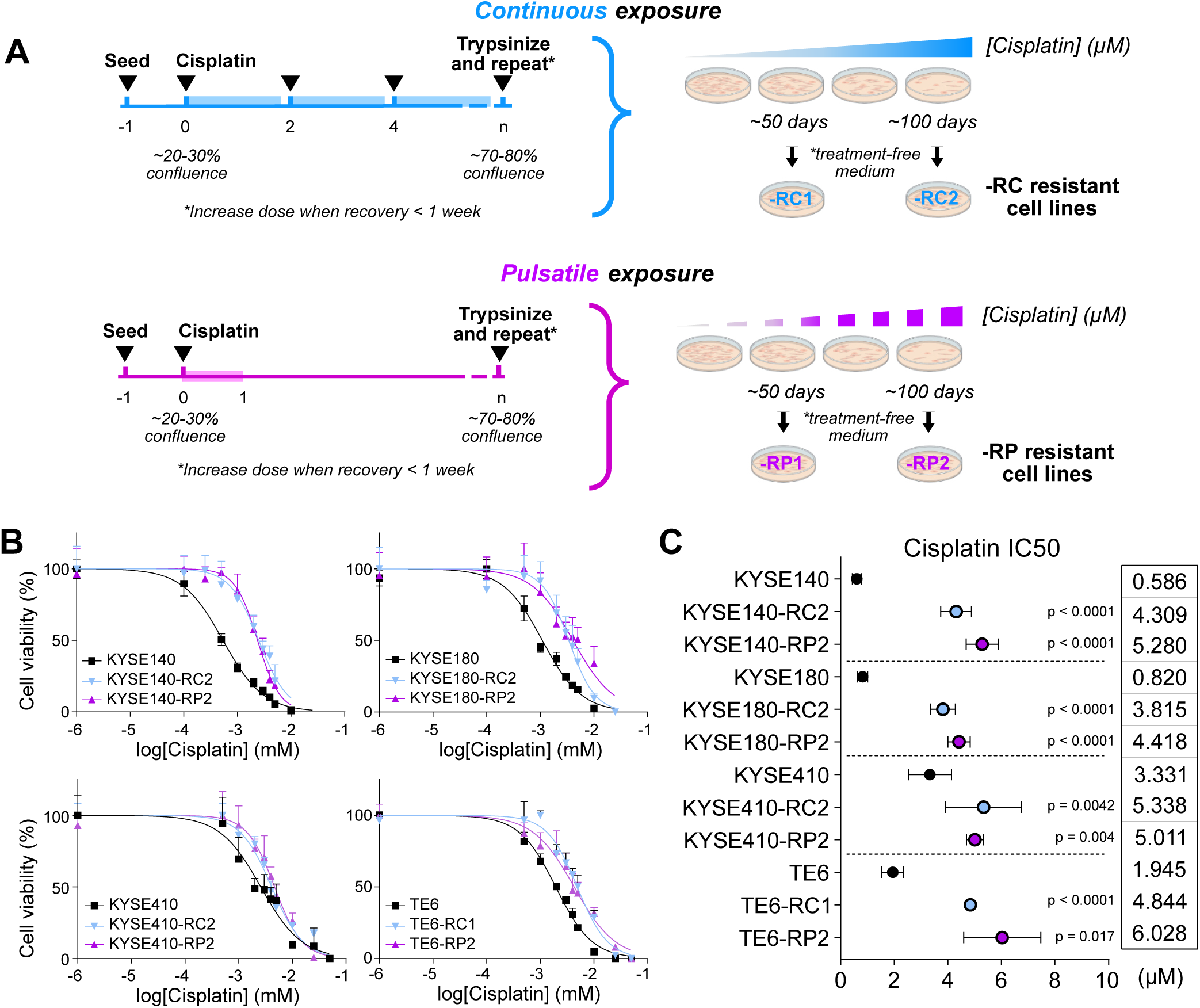
Cisplatin treatment results in chemotherapy resistance regardless of exposure regimen. **(A)** Experimental design for pulsatile vs continuous exposure. **(B)** Viability curves of KYSE140, KYSE180, KYSE410 and TE6 and their corresponding resistant clones. Cells were treated with increasing concentrations of cisplatin for 72h. Viability was measured using an MTT assay and normalized to vehicle control (0.9% NaCl). Data are mean ± SD (n = 6). **(C)** Summary of cisplatin IC50 values of parental and resistant cell lines. Data are mean ± SD (n = 3). Statistical significance of resistant versus parental cell lines was assessed with one-way ANOVA and Dunnett’s correction.

**Figure 2.**
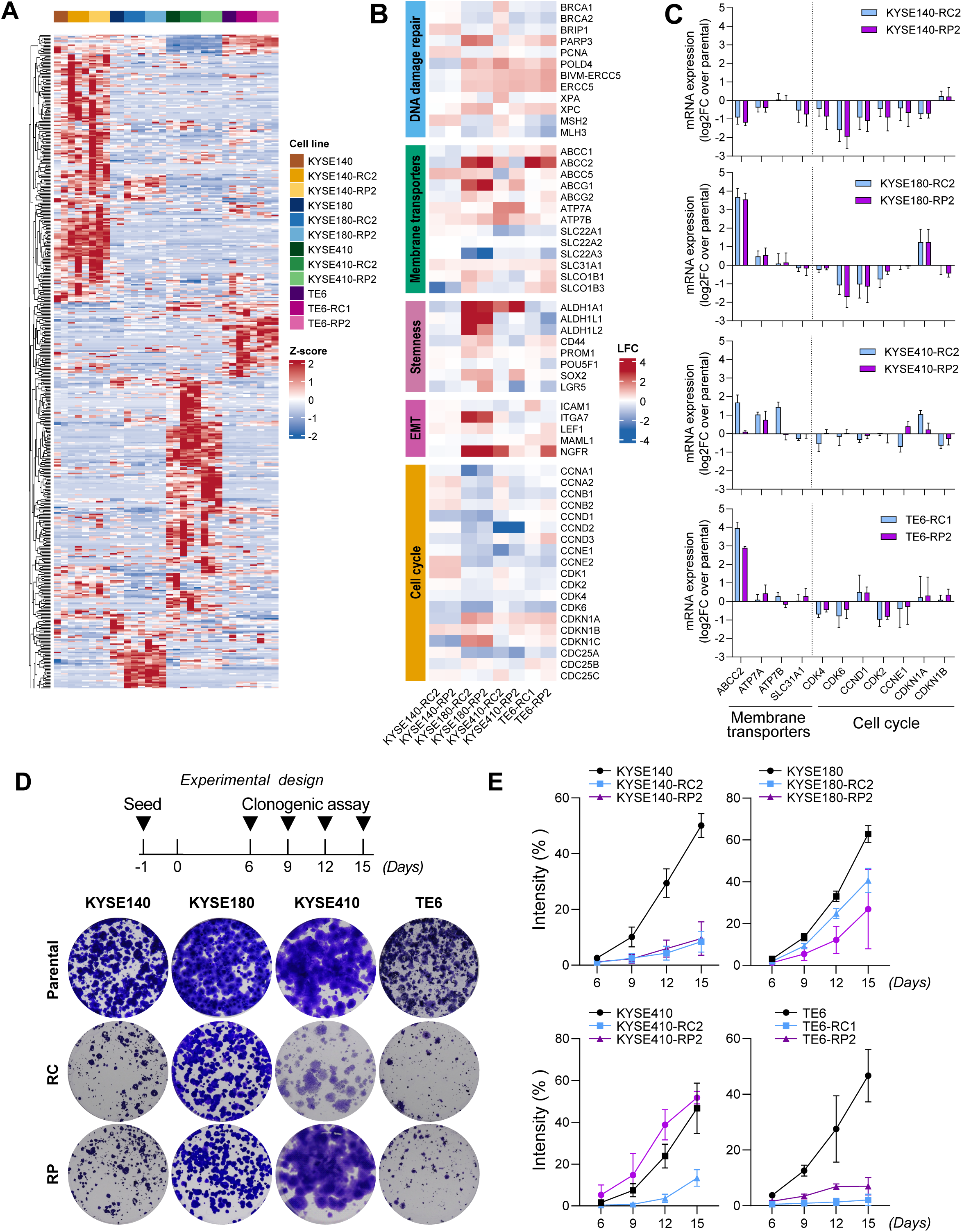
Cisplatin resistance induces transcriptional re-programming resulting in diverse adaptive mechanisms. **(A)** Z-score heatmap of RNAseq data from parental and resistant cell lines. **(B)** RNA-seq heatmap of resistant versus parental cell lines, showing significant differentially expressed genes related to common mechanisms of cisplatin resistance (DNA damage repair, membrane transporters, stemness markers and EMT markers) and the cell cycle. **(C)** qPCR validation of selected membrane transporters and cell cycle genes. Expression levels are shown as log2-fold change (log2FC) relative to parental. Data are mean ± SD of independent biological replicates (n = 3). **(D)** Experimental design of clonogenic assay with representative images of cell growth at day 15. **(E)** Difference in growth curves overtime between parental and resistant cell lines represented by intensity (%) of staining per pixel, a parameter that considers both the well area covered by the colonies and the density of the colonies. Data are mean ± SD (n = 3).

Resistant clones displayed both canonical and non-canonical resistance features. Classical mechanisms were observed in all clones, including the modulation of DNA damage repair pathways, such as the upregulation of NER-associated genes (*EERC5, POLD4, XPA, XPC*); or the modulation of membrane transporters, such as the upregulation of efflux transporters (*ABCC2/*MRP2) or downregulation of influx transporters (*SLC31A1*/CTR1, *SLCO1B3*/OATP1B3) (**Fig. 2B**), later confirmed by qPCR analyses (**Fig. 2C**). However, these changes occurred within a broader transcriptional reprogramming (**Fig. 2A**). Transcriptomic analyses revealed a general downregulation of cell cycle associated genes across most resistant clones (**Fig. 2B, C; Supplementary Fig. 1D**), suggesting a shift toward a low-proliferative state, confirmed by the reduced clonogenic capacity and slower growth kinetics (**Fig. 2D, E**). Concomitantly, most resistant cell lines exhibited upregulation of stemness-associated markers, *SOX2* and *ALDH1*; upregulation of epithelial-mesenchymal (EMT) markers, *NGFR* (CD271) and *MAML1*(**Fig. 2B**); and MYC activity suppression (**Supplementary Fig. 1E**), consistent with a drug-tolerant persister (DTP)-like phenotype. DTP cells are usually characterized by their reversibility, being transiently induced during treatment, but returning to the original state after drug withdrawal^13–15^. Interestingly, our ESCC resistant clones maintained a DTP-like state even in the absence of cisplatin (**Fig. 1A**), suggesting that cancer cells may sustain this cisplatin-induced, low-cycling state following drug removal in the absence of other external stressors, closer to a cancer stem cell-like phenotype^16–18^.

In contrast, KYSE410-RP2 underwent limited transcriptional reprogramming, retaining a transcriptional profile closely related to its parental (**Fig. 2A; Supplementary Fig. 1C**). Functionally, this was reflected in maintained proliferative capacity with no reduction in clonogenic growth (**Fig. 2D, E**). Consistent with this proliferative phenotype, KYSE410-RP2 lacked the mesenchymal and stemness markers observed in the other clones and showed instead upregulated *SOX2* expression (**Fig. 2B**), a transcription factor associated with enhanced proliferation and poor prognosis in squamous cell carcinomas^19^. This data suggested that KYSE410-RP2 underwent a different adaptive route than the other resistant clones, reinforcing its proliferative capacity rather than entering a quiescent state.

Together, these results establish that cisplatin exposure does not drive a binary resistant phenotype in ESCC but a continuum of adaptive cell states, ranging from quiescent persister-like cells to proliferative resistant populations. These data support a model in which chemotherapy exposure can promote state diversification, generating multiple adaptive routes to resistance from the same biological makeup.

### Cisplatin-induced plasticity drives cross-resistance to CDK4/6 inhibition

While we can expect cisplatin treatment to alter the molecular landscape of tumoral ESCC cells, the way it may modulate the response to subsequent therapy with CDK4/6i remains unknown. We previously reported that the CDK4/6 inhibitor palbociclib exerts both cytostatic and immunomodulatory effects in naïve ESCC cell lines^8^. Given that cisplatin exposure induced a broad transcriptional reprogramming of all resistant clones without a loss in Rb protein and phosphorylation levels (**Supplementary Fig. 2A**), we explored whether CDK4/6i, which acts through Rb-dependent cell cycle, would remain effective in cisplatin-resistant cells.

Treatment with the CDK4/6 inhibitors abemaciclib and palbociclib revealed a consistent shift in drug sensitivity across resistant clones, reflected by the increased EC50 values and enhanced proliferative capacity following treatment as compared to parental cells, especially in KYSE140 and KYSE180 resistant lines (**Fig. 3A-C; Supplementary Fig. 2B**). To better characterize the apparent loss of cytostatic effect following CDK4/6i, we treated all parental and resistant cell lines with a fixed concentration of palbociclib (1µM) for 15 days, or 6 days with a subsequent washout period to assess recovery (**Fig. 3D)**. KYSE140 and KYSE180 resistant clones did not only exhibit a significant increase in cell viability under continuous palbociclib exposure but also a significantly better recovery following drug withdrawal (**Fig. 3E-G; Supplementary Fig. 2C**). This enhanced recovery was particularly relevant, as it suggested that resistant cells were not merely tolerating CDK4/6i but actively retaining the capacity to resume proliferation once the selective pressure was removed, more consistent with a plastic, adaptable phenotype than with fixed genetic resistance.

**Figure 3.**
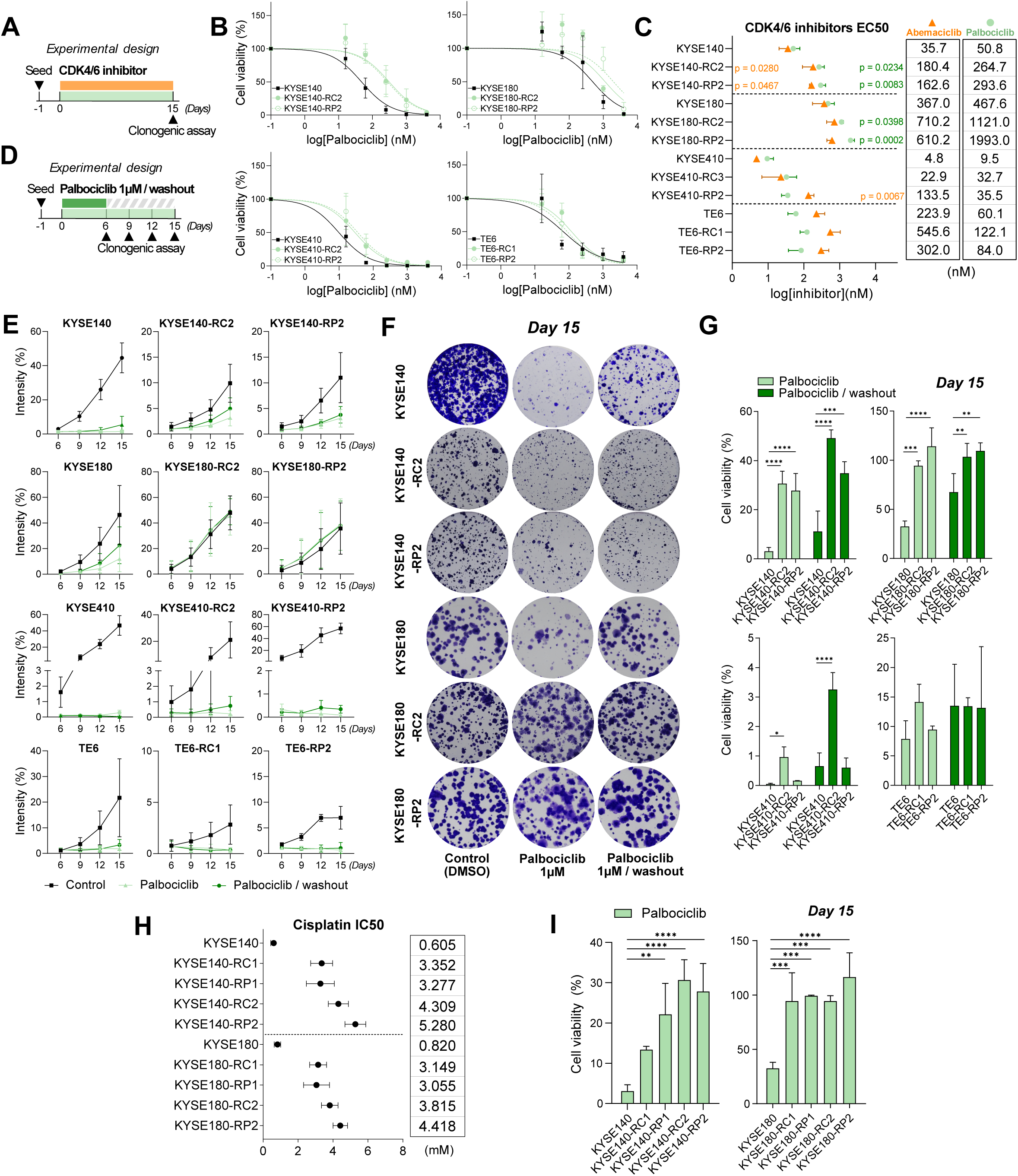
Cisplatin resistance drives loss of the cytotoxic effect associated to treatment with CDK4/6 inhibitors. **(A)** Experimental design to assess the EC50 of the CDK4/6 inhibitors abemaciclib and palbociclib. Cells were treated with increasing concentrations of abemaciclib (62-250-1000-2000-4000 nM) or palbociclib (16-62-250-1000-4000 nM) for 15 days prior to clonogenic assay. **(B)** Viability curves of KYSE140, KYSE180, KYSE410 and TE6 and their corresponding resistant clones treated with palbociclib as described in (A). Data are mean ± SD (n = 3). **(C)** Summary of abemaciclib and palbociclib EC50 values of parental and resistant cell lines. Data are mean ± SD (n = 3). **(D)** Experimental design to assess cell viability and cell recovery following palbociclib treatment. Parental and cisplatin-resistant ESCC cell lines were treated with 1μM palbociclib for 15 days, 6 days followed by a 9-day washout period, or vehicle control (DMSO). **(E)** Difference in growth curves overtime between parental and resistant cell lines as described in (D). Cell growth is represented by intensity of staining per pixel. Data are mean ± SD (n = 3). **(F)** Representative images at day 15 from (D). **(G)** Differences in cell viability (%) at day 15 from (D). The percentage of cell viability is derived from the formula: (%) = [(treated intensity / control intensity) * 100]. Data are mean ± SD (n = 3). Statistical significance was assessed with two-way Anova with Dunnett’s correction (∗ p < 0.05, ∗∗ p < 0.01, ∗∗∗ p < 0.001, ∗∗∗∗ p < 0.0001). **(H)** Summary of cisplatin IC50 values of KYSE140 and KYSE180 parental, early resistant (-R1) and late resistant (-R2) cell lines. Data are mean ± SD (n = 3). **(I)** Differences in cell viability (%) from cell lines were treated with 1 μM palbociclib for 15 days or vehicle control (DMSO). The percentage of cell viability is derived from the formula: (%) = [(treated intensity / control intensity) * 100]. Data are mean ± SD (n = 3). Statistical significance was assessed with one-way Anova with Dunnett’s multiple comparisons test (∗∗ p < 0.01, ∗∗∗ p < 0.001, ∗∗∗∗ p < 0.0001). Data for parental and -RC2/RP2 cell lines correspond to data presented in Fig. 1C, 3G, presented again for comparison with -RC1/RP1 lines.

We next investigated whether the depth of the cisplatin-induced reprogramming could determine the degree of cross-resistance to CDK4/6i by examining the early resistant clones (R1). These clones had been exposed to cisplatin for a shorter period, reaching lower IC50 values than their late resistant R2 counterparts (**Fig. 3H**). Shorter exposure to the platinum salt may have resulted in a lesser state of plasticity and, therefore, should be reflected on the cytostatic efficacy of palbociclib. Clonogenic assays (**Fig. 3I; Supplementary Fig. 2D**) showed that KYSE180-RC1/RP1 had already been driven into an almost complete loss of efficacy of the CDK4/6 inhibitor, analogous to the RC2/RP2 clones. However, KYSE140-RC1/RP1 clones demonstrated an intermediate state of sensitivity between the parental and the late counterparts (Fig. 15B; Sup. Fig. 2D), supporting the hypothesis of a continuum of plasticity concomitant with increased drug resistance.

Together, these data suggest that cisplatin-induced plasticity, which encompasses a continuum of resistant states, not only drives resistance to chemotherapy but also modulates the degree of cross-resistance to CDK4/6i in a state-dependent manner: the further cells are pushed along the cisplatin-induced plasticity continuum, the deeper and more entrenched their resistance to subsequent CDK4/6 inhibition becomes. Rather than an incidental consequence of platinum exposure, cross-resistance emerges as a predictable feature of the plastic state cisplatin induces.

### Multi-omic profiling links cisplatin-induced reprogramming to attenuated CDK4/6 inhibitor inflammatory response and emergent immune-evasive states

Our data established that cisplatin-induced plasticity drives cross-resistance to CDK4/6 inhibition in a depth-dependent manner. To better characterize the mechanisms underlying CDK4/6i cross-resistance, we performed integrative multi-omic analyses combining transcriptomic, proteomic, and phosphoproteomic analyses of parental and resistant clones following six days of palbociclib (1µM) treatment (**Fig. 4A**).

**Figure 4.**
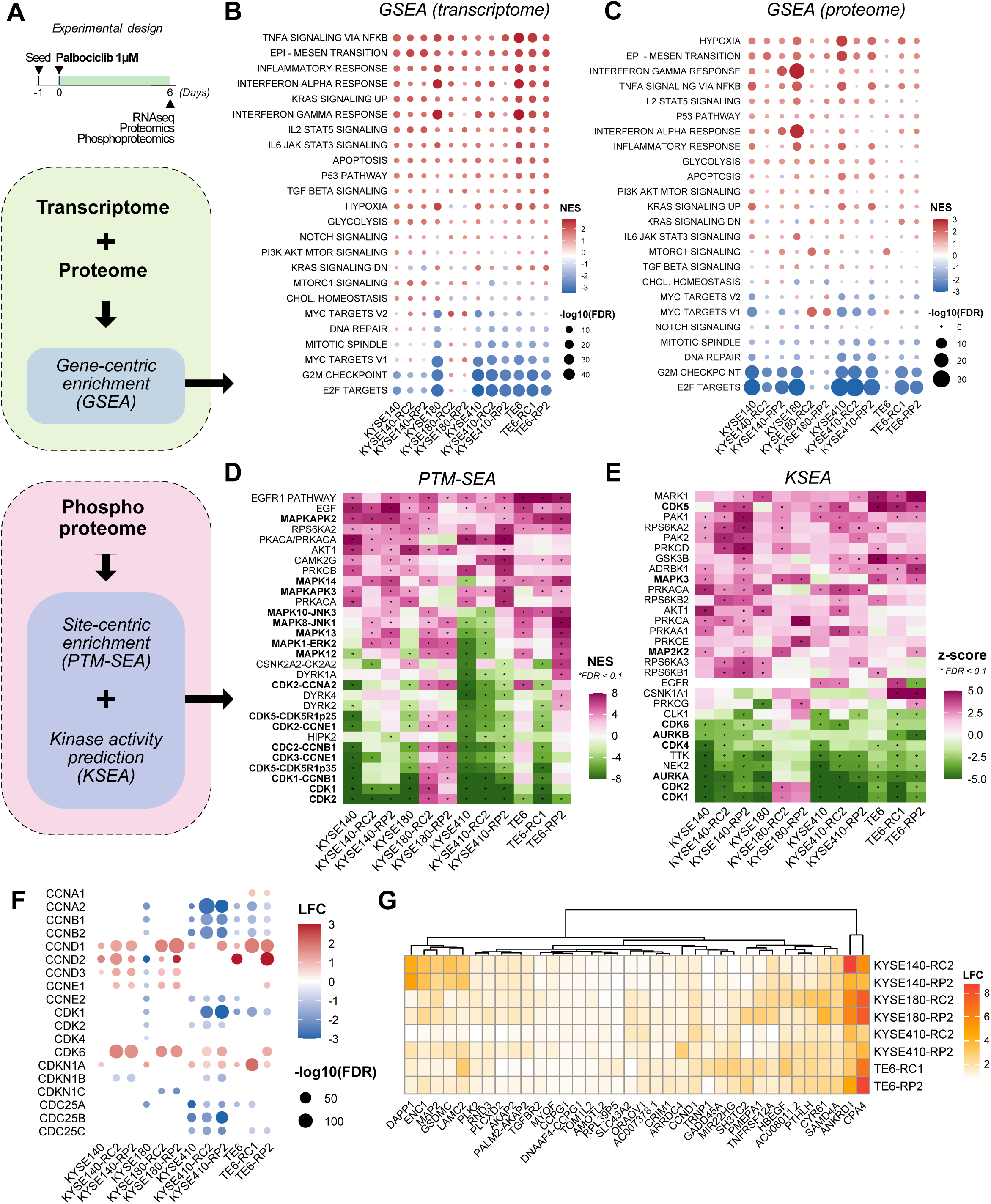
Multi-omic profiling reveals attenuation of CDK4/6 inhibition inflammatory response associated with emergent immune-evasive states in cisplatin-resistant clones. **(A)** Experimental design for the acquisition and analysis of multiomics data. Cell lines were treated with 1μM palbociclib for 6 days or vehicle control (DMSO), followed by transcriptome and (phospho)proteome acquisition. **(B)** Gene set enrichment analysis (GSEA) against Hallmark curated gene dataset from RNAseq data and **(C)** proteome data. **(D)** Heatmap of top 30 altered pathways from site-centric enrichment analysis (PTM-SEA) from phosphoproteome data. **(E)** Heatmap of top 30 kinases with altered activity predicted by kinase-susbtrate enrichment analysis (KSEA) from phosphoproteome data. **(F)** RNA-seq dotplot of cell lines treated with palbociclib 1μM for 6 days versus vehicle control (DMSO), showing significant differentially expressed genes related to cell cycle. Significance threshold was |LFC| > 0.585 and FDR < 0.1. **(G)** Heatmap of significantly upregulated DEGs during palbociclib treatment shared by all resistant cell lines (n genes = 36). Significance threshold was |LFC| > 0.585 and FDR < 0.1.

Gene set enrichment analysis (GSEA) from transcriptome and proteome data confirmed the loss of cytostatic effect previously observed with clonogenic assays. Multiple cell cycle-associated gene sets, such as “*E2F targets*”, “*G2M checkpoint*”, “*Mitotic spindle*”, “DNA repair” and “*MYC targets v1/2*”, were noticeably downregulated in all parental cell lines, but minimally (or even upregulated) in the KYSE140 and KYSE180 resistant clones (**Fig. 4B, C**). Similarly, site-centric enrichment analysis (PTM-SEA) and kinase activity prediction analysis (KSEA) from phosphoproteome data reflected how multiple cell cycle-related kinases that were inhibited in the parental lines were less so (or activated) in KYSE140 and KYSE180 resistant clones, including CDK1/2/3 or Aurora Kinase A/B (**Fig. 4D, E**). Interestingly, these resistant clones presented the activation of multiple signaling pathways known to drive palbociclib resistance in multiple models supported by both transcriptional and phosphoproteomic signatures, including the activation of multiple MAPK kinases (**Fig. 4D-E; Supplementary Fig. 3A**) and the upregulation of PI3K-AKT-mTOR pathway (**Fig. 4B-C; Supplementary Fig. 3B**) ^21–23^. Notably, while KYSE140 and KYSE180 resistant clones exhibited *CDK6* and *CCNE1* upregulation/activation and *CDKN1A* (p21) downregulation (**Fig. 4D-F**), these resistance markers were not elevated at baseline **(Fig. 2B,C)**. Conversely, their expression was only induced upon palbociclib exposure, further supporting the notion that cisplatin-induced plasticity enables adaptive responses to subsequent therapies.

Beyond these canonical resistance pathways, multi-omic profiling revealed a striking attenuation of immune and inflammatory signaling across all resistant clones (**Fig. 4B, C**). CDK4/6i is known to induce not only cell cycle arrest but also a pro-inflammatory phenotype through interferon (IFNα, γ) and NFκB signaling, a response linked to immune modulation in breast cancer^24–26^ and ESCC, as we previously reported^8^. While parental cells expectedly exhibited robust activation of interferon pathway upon palbociclib treatment, this response was markedly attenuated in resistant clones (**Supplementary Fig. 3C-E**). In parallel, the “*TNF*α *signaling via NFkB pathway*” gene set was also downregulated, accompanied by the upregulation of *ANKRD1,* a modulator of inflammation through inhibition of NF-κB (**Fig. 4B, C, G; Supplementary Fig. 3F**). Interestingly, recent research showed that pre-existing aberrant IFN or IL-6/STAT3 activation predicted CDK4/6i resistance in breast cancer patients and correlated with blunted therapy-induced inflammatory responses, EMT and immune-evasive signatures^27–29^. This raised the question whether the resistant clones in our model had already acquired a similarly dysregulated inflammatory baseline, which could explain the loss inflammatory signaling upon palbociclib exposure. Notably, transcriptional profiles from resistant clones prior to CDK4/6i revealed that most had already adopted this aberrant inflammatory state (**Supplementary Fig. 3G**). Consistent with an immune evasive phenotype, shared transcriptional signatures of palbociclib-treated clones showed upregulation of *CPA4* and *PMEPA1* (**Fig. 4G; Supplementary Fig. 3F**), two markers associated with tumor progression, drug resistance and immune exhaustion^30–35^.

Our data suggests that cisplatin resistant states are not defined by a single dominant pathway but instead exhibit a coordinated rewiring of signaling pathways, triggering a dysregulated inflammatory state that leads to immune-evasive phenotypes, which may contribute to CDK4/6 resistance in ESCC.

### Signature-based approaches identify HDAC inhibition as a strategy to reverse resistant cell states

Our data revealed that cisplatin resistance is not driven by a single dominant pathway but rather by a complex reprogramming of the transcriptional landscape. Therefore, rather than targeting individual markers, we aimed to identify compounds capable of broadly reverting the reprogrammed resistant states back toward the initial, therapy-sensitive state.

To this end, we leveraged the Connectivity Map (CMap) database, which comprises over ∼1.5 million datasets of transcriptional signatures of different cancer cell lines in response to thousands of treatments (**Fig. 5A**). In order to revert cisplatin resistance, we selected the significantly upregulated genes across all cisplatin-resistant cells (**Fig. 5B**), leading to a transcriptional signature of 36 genes (**Fig. 5C**), and queried it against CMap to look for the most dissimilar signatures. These signatures should oppose our signature and, therefore, be able to direct the resistant clones towards a more sensitive phenotype. Among the top candidates we found histone deacetylase (HDAC) inhibitors, glycogen synthase kinase-3 beta (GSK3β) inhibitors, and polo-like kinase (PLK) inhibitors (**Fig. 5D**). Interestingly, the top 5% compounds also included multiple tubulin inhibitors (e.g. vinorelbine, albendazole, flubendazole) associated to reversal of chemotherapy resistance in different cancers^36–38^.

**Figure 5.**
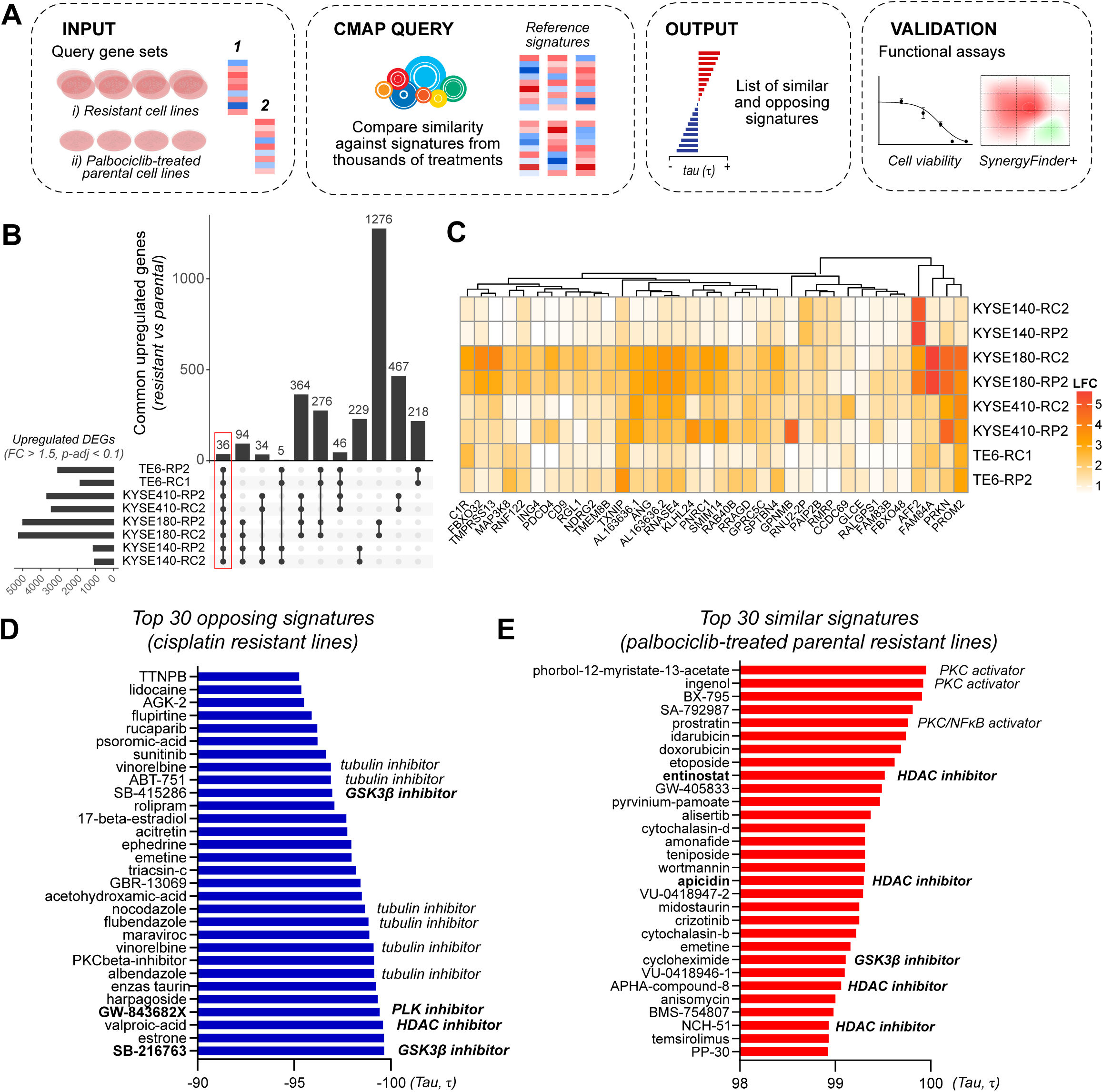
Signature-based approaches identify HDAC inhibition as a strategy to reverse resistant cell states. **(A)** Diagram of signature-based pipeline for drug candidate search and functional validation. Two signatures, (i) upregulated genes shared across all resistant clones (FC > 1.5, p-adj < 0.1), and (ii) upregulated genes shared across all parental cell lines following palbociclib treatment (FC > 1.5, p-adj < 0.1), were queried separately against CMap database. The output, two lists of similar and opposing drug signatures, were used for drug candidate selection followed by functional validation. **(B)** Upset plot from RNAseq data of significantly upregulated DEGs across resistant cell lines versus their parental, and **(C)** heatmap of transcriptional signature (i) shared by all resistant cell lines. Significance threshold was defined as FC > 1.5 and p-adj < 0.1. **(D)** Output from CMap query with the top 30 opposing drug signatures against signature (i), ranked by their connectivity score (tau). **(E)** Output from CMap query with the top 30 similar drug signatures against signature (ii), ranked by their connectivity score (tau).

A similar approach was applied to the transcriptional signature of the parental cell lines in response to palbociclib to predict whether any of these compounds could synergize with palbociclib to challenge the cross-resistance (**Supplementary Fig. 4A, B**). Notably, this second query further highlighted HDAC and GSK3β inhibitors as compounds capable of recapitulating key aspects of the parental transcriptional response (**Fig. 5E**). Interestingly, other top candidates included several PKC activators and a NFκB activator, demonstrating that the selected parental signature effectively recapitulated the pro-inflammatory nature of palbociclib. Moreover, this signature was clearly lost in the resistant cell lines (**Supplementary Fig. 4B**) in alignment with the results of the previous section.

In summary, these findings highlight HDAC, GSK3β and PLK inhibitors as potential candidates for the state-reversal strategy, aiming at reprogramming resistant cells toward a more therapy-sensitive state and regaining palbociclib efficacy. Together with the existing literature, the following candidates were selected for further study: apicidin, entinostat and vorinostat (HDAC inhibitors), SB-216763 (GSK3β inhibitor) and GW-843682X (PLK inhibitor).

### HDAC inhibition re-sensitizes resistant cells to cisplatin by reversing the resistant state

To validate whether all selected inhibitors could revert the resistant state, we assessed the cytotoxic effect of each inhibitor after 6 days of treatment, alone or in combination with cisplatin. Given the marked cytostatic resistance to palbociclib exhibited by KYSE140 and KYSE180 resistant clones, KYSE140-RP2 and KYSE180-RP2 were selected as representative cell lines for further investigation (**Fig. 6A**).

**Figure 6.**
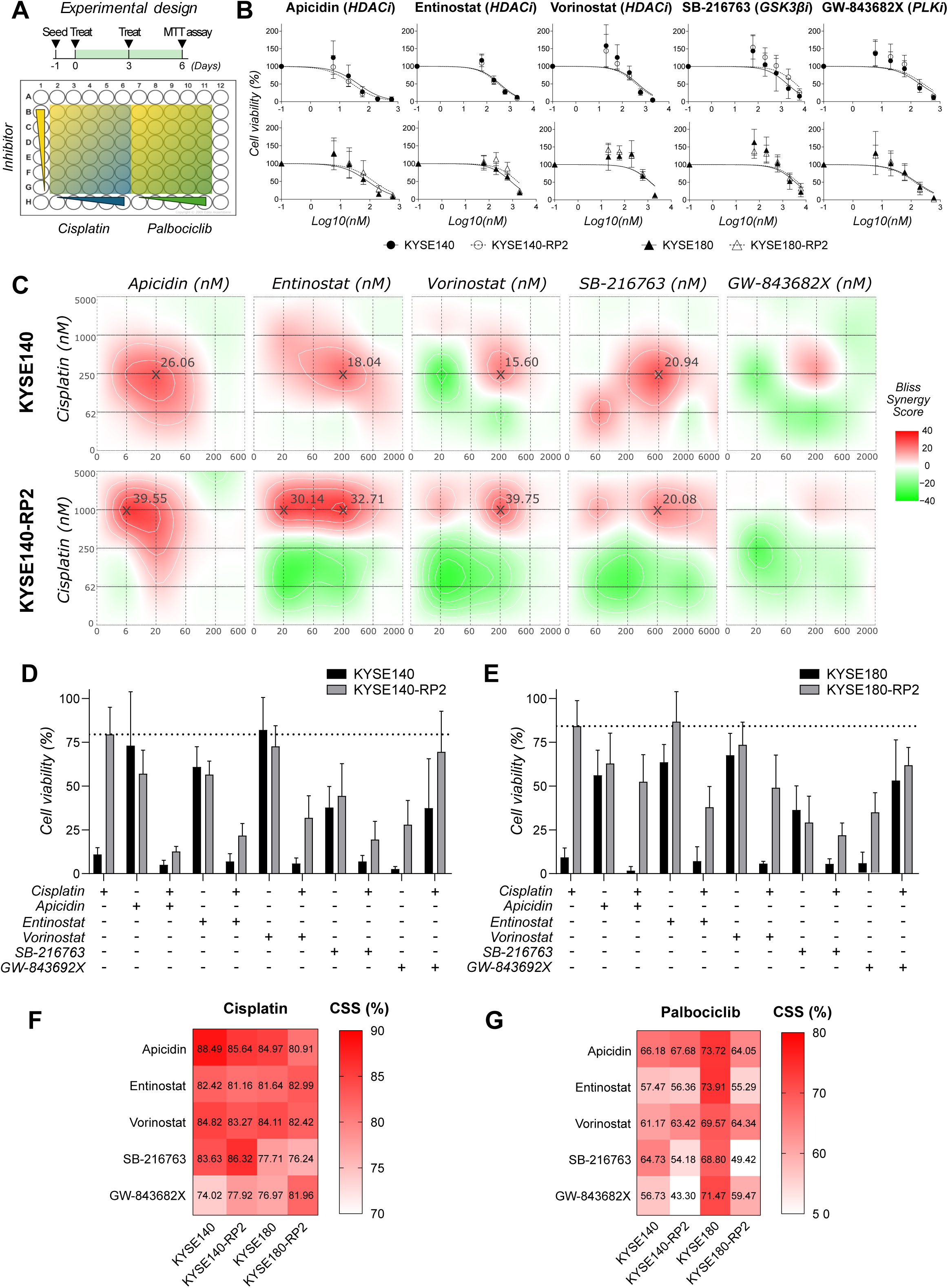
HDAC inhibition re-sensitizes resistant cells to cisplatin by reversing the resistant state. **(A)** Experimental design of synergy assay. Cells were treated with a combination of each inhibitor and cisplatin or palbociclib for 6 days (with a refresh at day 3) followed by MTT assay and SynergyFinder+ analysis. **(B)** Viability curves of KYSE140 and KYSE140-RP2 cell lines after treatment with monotherapy of signature-based selected inhibitors. Viability was normalized to vehicle control (DMSO). Data are mean ± SD (n = 3). **(C)** Bliss synergy scores heatmaps of KYSE140 and KYSE140-RP2 cell lines following combination of cisplatin with signature-based selected inhibitors. Synergy scores were calculated with (next page) SynergyFinder+ from 3 independent biological replicates (n = 3). Synergy scores above 10 were considered synergistic, scores between 0 and 10 additive, and scores below 0 antagonistic. **(D)** Bar plot of KYSE140 and KYSE140-RP2 cell lines for selected cisplatin-inhibitor combinations, with concentrations ranging cisplatin 1µM, apicidin 20 nM, entinostat 200 nM, vorinostat 200 nM, SB-216763 2000 nM and GW-843682X 200 nM, and **(E)** KYSE180 and KYSE180-RP2 cell lines, with concentrations ranging cisplatin 1µM, apicidin 60 nM, entinostat 600 nM, vorinostat 600 nM, SB-216763 2000 nM and GW-843682X 200 nM. Data are mean ± SD (n = 3). **(F)** Combination Sensitivity Scores (CSS) for combinations of HDAC inhibitors (apicidin, entinostat, vorinostat), GSK3β inhibitor (SB-216763) and PLK inhibitor (GW-843682X) with cisplatin and **(G)** palbociclib. CSS scores were calculated with SynergyFinder+ from 3 independent biological replicates (n = 3). A CSS > 50% indicates that the drug combination achieved greater than 50% maximal cell inhibition over the tested concentration range. Combinations with a CSS > 75% were considered exceptionally sensitivity.

Viability curves from monotherapy with each inhibitor showed that most candidates did not experience cross-resistance, with similar IC50 values between parental and resistant clones (**Fig. 6B**). SynergyFinder+^39^ was then used to calculate the synergy score of combined therapies. All HDAC inhibitors (apicidin, entinostat, vorinostat) and the GSK3β inhibitor (SB-216763) consistently demonstrated strong synergistic effects with cisplatin (**Fig. 6C; Supplementary Fig. 5A**), restoring sensitivity in resistant clones and, in some cases, fully reverting the resistant phenotype to parental levels (**Fig. 6D, E**). Notably, all combinations were associated high combination sensitivity scores (CSS > 75%) (**Fig. 6F**). While synergy is relevant for minimising drug doses and reducing systemic toxicity, a high CSS (i.e. efficacy of the drug combination at killing or inhibiting the cells) is necessary for effective treatment. Interestingly, HDAC inhibitors also demonstrated high synergy and CSS with cisplatin in the parental cell lines, suggesting that cisplatin may benefit from combination with HDAC inhibitors regardless of the resistant status (**Fig. 6C, F; Supplementary Fig. 5A**). Together, these data support a model in which HDAC inhibition acts as a reprogramming agent not only capable of reverting the resistant transcriptional state, but also of acting as cytotoxic enhancer in combination with cisplatin prior to establishment of acquired resistance.

We also investigated whether reversing the cisplatin-resistant state could also restore sensitivity to CDK4/6i. Combination treatments with HDAC inhibitors and palbociclib showed variable but measurable improvements in response, depending on the cellular context and degree of prior reprogramming (**Fig. 6G**). While synergy was less pronounced than with cisplatin (**Supplementary Fig. 5B**), these results suggest that state reprogramming could partially overcome CDK4/6i cross-resistance.

Together, our findings support a model in which cisplatin exposure drives cancer cells into a dynamic landscape of adaptive states characterized by transcriptional plasticity, phenotypic diversification, and emergent vulnerabilities. Within this framework, resistance can be intercepted through state-reprogramming strategies, such as HDAC inhibition, enabling rational combination therapies aimed at restoring drug sensitivity and preventing cross-resistance.

## Discussion

Despite research efforts, esophageal squamous cell carcinoma (ESCC) remains a disease with limited therapeutic options and poor long-term outcomes. Platinum-based regimens remain the standard of care in this setting, but 5-year survival rates are as low as 20% and most patients relapse^40–43^. The capacity of tumor cells to adapt under treatment pressure through genetic modifications, transcriptional plasticity and interactions with the tumor microenvironment collectively contribute to the emergence of acquired resistance. A better understanding of these mechanisms may help identify new opportunities for therapeutic intervention.

### Cisplatin induces a dynamic landscape of adaptive resistant states

Acquired resistance to chemotherapy is increasingly understood as a dynamic process modulated by both genetic and non-genetic mechanisms rather than a single dominant alteration, giving rise to plastic phenotypic states. In line with this view, we found that cisplatin exposure in ESCC cells drove a broad transcriptional reprogramming leading to a spectrum of adaptive states with different proliferative behavior. Notably, most resistant clones showed traits consistent with a DTP phenotype. Since their discovery in 2010^44^, multiple studies have described the presence of DTPs in melanoma, lung, breast, colorectal, gastric, or head and neck cancer^45–50^, but only one study has addressed DTP behavior in ESCC cells^51^. All formal definitions of DTPs delimit these states as transient, reversible phenotypes that are resolved following drug withdrawal^14,15^. Interestingly, our cisplatin-resistant ESCC cells revealed a persistence of the slow-cycling and drug tolerant state in spite of the absence of drug pressure, and presented additional canonical mechanisms of platinum resistance. These results supported the theory proposed by a growing body of research suggesting that, under prolonged exposure, DTPs have the potential to stabilize and progress towards more advanced states of resistance, ultimately serving as a pool for cancer recurrence and refractory treatment^14,52^.

While it is clear that drug exposure is a driver of acquired drug resistance, prolonged treatment may act as a stochastic bottlenecking that limits the diversity of these adaptive, resistant states^53^. In this study, we showed that clones derived from the same parental line converged more strongly with one another than with clones from different parental backgrounds, suggesting that resistance plasticity is constrained by pre-existing lineage context and that exposure to the same drug may lead to converging mechanisms regardless of treatment regimen. However, similar to Ramirez *et al.* (2016), we also found that therapy exposure can also lead to phenotypic diversification^52^. Albeit KYSE410-RP2 and KYSE410-RC2 shared molecular strategies of cisplatin resistance, they showed divergent trajectories towards both a slow-cycling, DTP-like phenotype, and a highly proliferative, stem-like state. Notably, KYSE410 is the least sensitive cell line at baseline. Because of prior intrinsic mechanisms of resistance, pulsatile exposure could have resulted in suboptimal cytotoxic drug concentrations, allowing the resistant KYSE410-RP2 cell line to maintain a highly proliferative phenotype. Interestingly, these results modeled heterogeneous drug delivery within the tumor niche, where spatial constraints in drug penetration can lead to ineffective drug exposure in some subpopulations, under which highly proliferative resistant clones may survive^54,55^. Oppositely, the continuous setting (replicating an “optimal” drug delivery) negatively impacted highly proliferative cells and favored the emergence of the slow-cycling phenotype of KYSE410-RC2. In this context, it remains a question whether the slow-cycling resistant clones arise from the induction of new cellular states or selection of pre-existing subpopulations. Interestingly, several studies using lineage-tracing and single-cell longitudinal approaches attempted to distinguish these mechanisms and demonstrated that both could give rise to DTPs^49,56^. Regardless of the initiating mechanism, understanding how this continuum of phenotypic adaptation evolves over time and how different states of phenotypic plasticity translate into different degrees of therapy resistance remain an important question. Within this framework, the early resistant clones (-R1) generated in this study may be used as a model of progression of resistance to examine, in combination with RNAseq and orthogonal assays such as ATAC-seq or CUT&Tag for histone acetylation marks (H3K27ac, H3K9ac), how intermediate adaptive states have evolved along the resistance continuum and to decorticate which epigenetic cues may orchestrate the resistant transcriptional reprogramming.

### Cisplatin-driven reprogramming promotes cross-resistance to CDK4/6 inhibition through dual cytostatic and immune-regulatory mechanisms

The inherent plasticity present in DTP cells allows them to switch between multiple phenotypic states, providing them with the ability to adopt dynamic strategies of resistance^57–59^. In our study, we demonstrated that cisplatin-resistant cell lines exhibited cross resistance to CDK4/6 inhibitors with a marked loss of cytostatic response, an aspect that had not yet been addressed in ESCC. Moreover, while most studies correlated palbociclib resistance to stable modifications, such as genetic amplification of *CCNE1* and *CDK6*^60,61^ or constitutive overexpression of 3-phosphoinositide-dependent protein kinase 1 (PDK1)^62^, we found that our cisplatin-resistant cell lines only expressed these markers upon palbociclib exposure rather than constitutively. These data supported that it was transcriptional plasticity, rather than fixed mechanisms of resistance, that conferred the cisplatin-resistant cells with the capacity to adapt under palbociclib insults. Interestingly, despite the well-documented contribution of the inflammatory response to CDK4/6i efficacy^25,26,63–65^, the mechanisms of resistance that may hamper this response remain poorly understood. Among the few studies that have addressed the loss of the inflammatory response, De Angelis *et al.* (2021) found that aberrant IFN activation in pre-treated tumors predicted poor response and correlated with a lack of palbociclib-induced IFN response in breast cancer patients^27^. Eudald *et al.* (2025) also revealed that chronic IFN-γ/STAT1 activation prior to treatment was associated with an immunosuppressive microenvironment and predicted shorter PFS and OS in an independent neoadjuvant palbociclib trial (NeoPalAna)^66^. Similarly, we found that our ESCC cisplatin-resistant cell lines were characterized by a baseline aberrant immune activation which correlated with a hindered IFN pathway activation following palbociclib response. This finding carries direct clinical implications: it provides a mechanistic explanation for the limited clinical activity of CDK4/6i reported in several trials (NCT01037790, NCT03339843), where all enrolled ESCC patients had previously undergone chemotherapy as primary line treatment^67^. Together, our study underlines a novel connection, by which cisplatin resistance can drive the loss of both the cytostatic effect exerted by CDK4/6i and the immune-permissive mechanisms necessary for CDK4/6i response. However, our conclusions are based on cell-intrinsic models and cannot resolve how these resistant states interact with stromal or immune compartments. This is particularly relevant for our interpretation of palbociclib-associated inflammatory signaling, which should be validated in cocultured 3D systems^68^ or humanized mouse models^69^.

### Emergent vulnerabilities and therapeutic implications

Provided the transcriptional rewiring that DTP-like cells undergo following drug exposure, new treatment strategies should aim at state-wide reversal approaches rather than single molecular targets. In this study, we employed a signature-based strategy and identified HDAC inhibitors as potential candidates for reversal of the resistant transcriptional program. Given the fundamental role of HDACs in chromatin accessibility and transcriptional regulation, the signature-based identification of HDAC inhibitors as potential targets to revert resistance was consistent with our hypothesis of cellular plasticity as driver of cisplatin tolerance and resistance. Interestingly, multiple HDAC inhibitors have been investigated as possible targets to revert chemotherapy resistance in lymphoma, prostate, bladder, lung, ovarian and oral cancer^70–78^. Similarly, HDACi synergized with cisplatin in ESCC cisplatin-resistant cell lines, re-sensitizing them towards basal levels of cisplatin response, in line with Xiao-Ping et al. (2018) in ESCC cells and xenograft models^79^.

The recognition that cancer drug resistance is, predominantly, a dynamic and continuous process rather than a unique, fixed genetic state has great therapeutic implications. The temporal evolution of therapeutic resistance, from initial DTP survival through epigenetic consolidation to stable genetic resistance, delimits a series of windows of intervention in which different therapeutic strategies should be prioritized^53,55,56^. In this context, intervention may be most effective at earlier stages when cells still occupy lower plastic, drug-tolerant states, and before non-genetic tolerance has shaped the emergence of more stable resistance mechanisms. Interestingly, synergy was not restricted to the resistant clones: cisplatin and HDAC inhibitors also acted synergistically in the parental cell lines. In that framework, the benefit of HDACi may not only lie in the re-sensitization of already resistant populations, but also in the prevention of the development of cisplatin-driven resistance if delivered in an appropriate schedule in ESCC, a strategy with clear translational implications. Indeed, Singh *et al.* (2025) showed that sequential HDACi and cisplatin synergistically impeded DTP cell emergence and survival in gastric and liver cancer cells^80^, whereas Xu *et al.* (2025) reported that knockdown of HDAC2/6/9 markedly reduced the formation of DTP cells and that combination of a HDAC inhibitor (YFF-702) with 5-fluorouracil exerted synergistic cytotoxic effect in DTP cells and in an *in vivo* tumoral model of ESCC^51^. Furthermore, if HDAC inhibition can prevent the cisplatin-induced epigenetic state, it may also avert the development of the immune evasive phenotype contributing to CDK4/6i cross resistance and rescue its immunological efficacy. Notably, multiple studies have unveiled the capacity of HDACi to restore tumoral immunogenicity and modulate the immune microenvironment towards a “hot”, tumor-suppressive phenotype, including esophageal cancer ^81–85^. It will be interesting, therefore, to examine whether a three-agent sequential strategy, combining cisplatin with HDAC inhibition to intercept the resistant state and CDK4/6 inhibition to promote the immune program, may improve response and prevent the emergence of refractory disease in ESCC patients.

In summary, cisplatin exposure drives a dynamic landscape of adaptive states in ESCC, dominated by a DTP-like phenotype that persists beyond drug withdrawal and promotes cross-resistance to CDK4/6 inhibition through concurrent loss of cytostatic and immune-permissive mechanisms. These findings lay the foundation for therapeutic strategies aimed at intercepting epigenetic reprogramming before resistance consolidation, providing a rationale for sequential HDAC inhibitor-based combinations to enhance cisplatin and CDK4/6 inhibitor sensitivity in ESCC.

## Materials and Methods

### Cell culture

KYSE140, KYSE180, KYSE410 ESCC cell lines were obtained from DSMZ (Brunswick, Germany) and TE6 ESCC cell line was obtained from the cell bank of the RIKEN BRC (Tsukuba, Japan). All cells were cultured in RPMI medium 1640 1X (Gibco, #52400-025) supplemented with 1% penicillin/streptomycin (Gibco, #15070-063) and 10% fetal bovine serum (Gibco, #A5256701). Cells were maintained in a 5% CO2 atmosphere and constant humidity.

### Drugs and inhibitors

Cisplatin (#479306) was purchased from Sigma Aldrich and dissolved in 0.9% NaCl at a stock concentration of 0.6 mg/mL (2 mM). Abemaciclib (LY2835219, #S7158) and palbociclib (PD-0332991 hydrochloride, #S1116) were purchased from SelleckChem and dissolved in dimethyl sulfoxide (DMSO) at stock concentrations of 5 mM. Apicidin (#HY-N6735), Entinostat (#HY-12163), GW-843682X (#HY-11003), SB-216763 (#HY-12012), and Vorinostat (#HY-10221) were purchased from MedChemExpress and dissolved in DMSO at stock concentrations of 1 mM, 10 mM, 1 mM, 10 mM, and 10 mM, respectively. All stock solutions were stored according to the manufacturer’s recommendations and diluted in complete culture medium to the desired working concentrations immediately prior to treatment.

### Generation of cisplatin-resistant cell lines

KYSE140, KYSE180, KYSE410 and TE6 cell lines were seeded in 6-well plates at densities ranging from 5 × 10^4^ to 1 × 10D cells per well, to be treated the day after at a 20-30% confluence. For the continuous method (“-RC”), the first treatment corresponded to the individual cisplatin IC25 of each cell line and was refreshed every 2-3 days. For the pulsatile method (“-RP”), the first treatment corresponded to the individual cisplatin IC75 of each cell line and was removed after 24h. In both methods, cells were trypsinized after reaching 70-80% confluence and re-seeded in a new 6-well plate to start a next round of treatment. The same doses were maintained until cells reached 70-80% confluence in less than a week, moment after which the treatment dose was increased. Early resistant clones (“-RC1/RP1”) and late resistant clones (“-RC2/RP2”) were kept under treatment with each method for approximately 50 and 100 days, respectively. Resistant clones were then maintained in treatment-free RPMI 1640 1X medium for a month prior to the assessment of cisplatin half-maximal inhibitory concentration (IC50) and the establishment of stable resistance cell lines.

### Cell viability assay

Cells were seeded in 96-well plates at densities ranging from 1 × 10^3^ to 6 × 10^3^ cells per well in a final volume of 150 µL per well. To minimize edge-effect evaporation, the outer wells were filled with 200 µL PBS. After 24h, cells were treated with concentrations of cisplatin ranging 0.1-50 µM, with six technical replicates per concentration (n = 6), and incubated for 72 hours. Cell viability was assessed using the MTT (3-(4,5-dimethylthiazol-2-yl)-2,5-diphenyltetrazolium bromide; Sigma Aldrich, #M5655) assay. MTT reagent was added to each well at a concentration of 0.5mg/ml and plates were incubated for 1.5 hours. The supernatant was then carefully discarded, and 100 µL of DMSO were added to each well to solubilize the formazan crystals. Plates were incubated at room temperature (RT) for 30 minutes with gentle agitation and absorbance was measured at 570 nm. Absorbance values were blank-corrected, and IC50 values were calculated in GraphPad Prism v8 from dose–response curves fitted to a four-parameter nonlinear regression model (*log(inhibitor) vs. normalized response -- variable slope*). Cell viability assays were performed at least in triplicate. IC50 values are reported as mean ± SD, and differences in IC50 values across cell lines were assessed by one-way ANOVA with Dunnett’s multiple comparisons test (p < 0.05).

### Clonogenic assay

Cells were seeded in 6-well plates at densities ranging from 2 × 10² to 4 × 10³ cells per well. To assess the differences in proliferative capacity between parental and resistant cell lines, each parental and its derived resistant clones were seeded with the same density and maintained in culture for 15 days without treatment. To determine the half-maximal effective concentration (EC50) of abemaciclib and palbociclib, cells were treated with either control (DMSO) or increasing drug concentrations ranging from 16 to 4000 nM for 15 days. To characterize drug response and recovery dynamics, cells were treated with either DMSO or 1 µM palbociclib for the full 15-day period (response condition), or for 6 days followed by a washout period with fresh medium until day 15 (recovery condition). In all conditions, culture medium and treatment were refreshed every 2 days. At each predetermined time point, cells were washed with PBS and fixed in 10% neutral buffered formalin (Sigma-Aldrich, #HT501128-4L) for 10 minutes at RT. Plates were then stained with 0.05% crystal violet (Sigma-Aldrich, #548-62-9) for 30 minutes under slow agitation, washed thoroughly with water and air-dried. All experiments were performed in triplicate. Plates were imaged with an Olympus PEN E-PL8 camera, and covered surface pixel intensity was quantified using the ColonyArea plugin in ImageJ. Intensity values were normalized relative to the control using the following formula: Cell viability (%) = [(intensity {treatment} / intensity {control}) * 100]. EC50 values were calculated in GraphPad Prism v8 from dose–response curves fitted to a three-parameter nonlinear regression model with standard slope (*log(inhibitor) vs. normalized response*) due to limited data points as per GraphPad guidelines. Exceptionally, the dose–response curve of parental TE6 cell line treated with palbociclib was fitted to a four-parameter nonlinear regression model (*log(inhibitor) vs. normalized response -- variable slope*), as it yielded a superior goodness of fit and captured the Hill slope more accurately. All assays were repeated in triplicate. Data are presented as mean ± SD. Statistical differences in EC50 of resistant versus parental clones were assessed by one-way ANOVA. Statistical differences between resistant and parental clones treated with palbociclib 1 µM with or without washout were assessed by two-way ANOVA. Multiple comparisons correction was performed with Dunnett’s test. Significance was denoted as *p < 0.05, **p < 0.01, ***p < 0.001, ****p < 0.0001.

### Immunoblotting

Cells were seeded in 60 mm dishes at densities ranging from 3 × 10D to 6 × 10D cells per dish and cultured until reaching 80% confluence. Cells were washed twice with ice-cold PBS and lysed by scraping in ice-cold Laemmli buffer supplemented with protease and phosphatase inhibitors (4 mM NaF, 100 µM sodium vanadate, 60 µg/mL Pefabloc SC, 1 µg/mL leupeptin). Lysates were transferred to pre-cooled microcentrifuge tubes and sonicated with a tip sonicator at 25% amplitude, 5-second ON/OFF cycles for a total of 45–60 seconds. Samples were denatured at 95°C for 5 minutes and centrifuged at 16,000 × g for 15 minutes at 4°C to remove insoluble debris. Clarified supernatants were stored at −80°C until further use. Protein concentration was determined by filter paper assay^86^. Equal amounts of protein (10 µg per sample) were separated by SDS-PAGE on 9% polyacrylamide gels and transferred onto Immobilon®-P PVDF membranes (0.45 µm, Millipore, #IPHV00010) by overnight wet transfer at 26 V, 4°C. Membranes were blocked for 1 hour RT in blocking buffer (5% non-fat dry milk {Bio-Rad #1706404}, 0.1% Tween-20 in TBS) and subsequently incubated with primary antibodies diluted in blocking buffer overnight at 4°C with gentle agitation. Membranes were washed three times for 15 minutes in blocking buffer and incubated with the appropriate HRP-conjugated secondary antibody (1:2000 in blocking buffer) for 1 hour at RT with gentle agitation. Membranes were then washed sequentially with blocking buffer, TBS-Tween, and TBS (15 minutes each). Protein signal was detected by enhanced chemiluminescence using Western Lightning® Plus-ECL (PerkinElmer, #NEL104001EA) and imaged on a Fusion solo S (Vilber Lourmat) optical system. All primary and secondary antibodies are listed in **Table 1**.

**Table 1.** List of antibodies for immunoblotting.

| Antibody | Manufacturer | Cat. No | Origin | Dilution |
| --- | --- | --- | --- | --- |
| **p-RB (Ser807/Thr811) XP** | Cell Signaling Technology | 8516 | Rabbit | 1:1000 |
| **RB** | BD Pharmingen | 554136 | Mouse | 1:500 |
| **HSP90** | BD Pharmingen | 610419 | Mouse | 1:500 |
| **Anti-mouse IgG, HRP-linked Antibody** | Cell Signaling Technology | 7076 | Horse | 1:2000 |
| **Anti-rabbit IgG, HRP-linked Antibody** | Cell Signaling Technology | 7074 | Goat | 1:2000 |

### Quantitative RT-PCR

Cells were seeded in 60 mm dishes at densities ranging from 3 × 10D to 6 × 10D cells per dish and cultured until reaching 80% confluence. Cells were washed twice with ice-cold PBS and lysed by scraping in TRK lysis buffer. Total RNA was isolated using E.Z.N.A.® MicroElute® Total RNA Kit (Omega bio-tek, # R6831), and 1µg RNA per sample was retrotranscribed with RNA Superscript II Reverse Transcriptase (Invitrogen, #18064). All conditions were obtained in biological triplicates. Quantitative RT-PCR was performed with 2 ng of cDNA per sample and KAPA SYBR® FAST qPCR Master Mix (2X) Kit (Kapa Biosystems, #KK4601) in a QuantStudio 3 real-time PCR system (Applied Biosystems) according to manufacturer’s protocols. All samples were analyzed in technical duplicate and *eEF2* gene was used for normalization. Primers were designed using the Integrated DNA Technologies primer designing tool(https://eu.idtdna.com/scitools/Applications/RealTimePCR/Default.aspx) and are listed in **Table 2**. Quantitative RT-PCR analysis was performed using the ΔΔCt method. Data are presented as mean + SD.

**Table 2.**
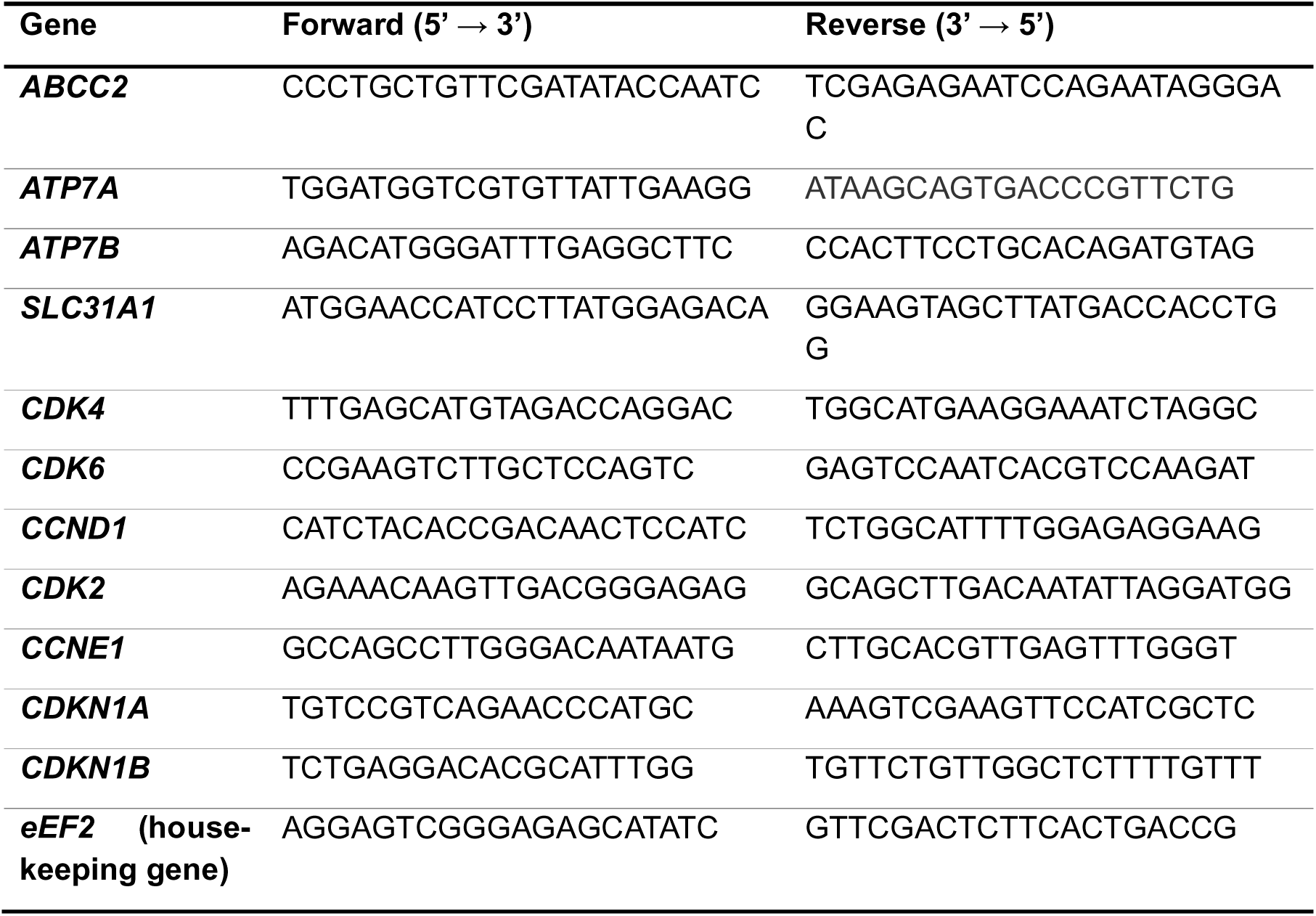
List of primers for qPCR.

### RNA-sequencing

#### Sample preparation and sequencing

Resistant cell lines were were seeded in 60 mm dishes at densities ranging from 3 × 10D to 6 × 10D cells per dish and treated with 1 µM palbociclib or vehicle control (DMSO) for 6 days. Total RNA was isolated using E.Z.N.A.® MicroElute® Total RNA Kit according to manufacturer’s protocol. RNA quality was assessed using a Fragment Analyzer 5200 (Agilent Technologies). One hundred nanograms of total RNA were used to construct indexed cDNA libraries using the NEBNext Ultra II Directional RNA Library Prep Kit for Illumina (New England Biolabs) according to manufacturer’s instructions. The multiplexed libraries were sequenced on an Illumina NovaSeq 6000 system using a 200 Cycles S2 flow cell at the BRIGHTcore facilty (Brussels, Belgium).

#### Data analysis

Paired-end reads were mapped against the human reference genome GRCm38 using STAR software (version 2.5.3a) and transcript annotations were obtained from Homo_sapiens.GRCh38.90.gtf (ftp.Ensembl.org). Gene-level counts were generated using HTSeq and normalized to counts per 20 million reads (CP20M). Raw data from parental (GEO accession: GSE312063) and from resistant cell lines were merged into a unique dataframe for analysis. Data quality was assessed in rlog-transformed data with Pearson’s correlation and principal component analysis (PCA) prior to analysis. Genes with low expression were filtered, retaining those with > 0.5 CPM in at least two samples. Differential expression analysis (DEA) was performed with with DESeq2 package (v1.40.2), and log2 fold-change (log2FC) values were shrunk with the lfcShrink function from the apeglm package (v1.24.0) to reduce noise from lowly expressed genes^87^. Significance cut-off was set at adjusted p-value < 0.1 and |LFC| > 0.585. Heatmaps were represented using the pheatmap package (v. 1.0.13), and hierarchical clustering was performed using Euclidean distance and complete linkage. Gene set enrichment analysis (GSEA) was performed using the fgsea package (v. 1.34.2). Genes were ranked by Log2FC and enriched against curated MSigDB gene set collections, including Hallmark and KEGG. The fgseaMultilevel() function was used with parameters minSize = 10 and maxSize = 1000. Enrichment scores were computed as normalized enrichment scores (NES), and significance was assessed by permutation testing. All analyses were performed with R (v 4.4.3).

### (Phospho)proteome extraction, LC-MS/MS acquisition and analysis

#### Sample preparation

Cells were seeded in 10cm dishes at 1.5 – 5.5 × 105 cells per dish overnight and treated with either vehicle control (DMSO) or palbociclib (1 µM) for 6 days, refreshing the treatment every 2 days. Prior to lysis, cells were incubated with freshly prepared pervanadate 0.1mM, washed twice with ice-cold PBS and once with ice-cold milliQ water. Samples were scratched in 500ul freshly prepared lysis buffer (8M urea, 50mM Tris-HCl pH 8.5, 1mM β-glycerolphosphate, 1mM NaF), sonicated at 26% amplitude, 5 s ON / 5 s OFF for 1 min and centrifuged at 16000 g for 15 min. Supernatants were stored at −80D°C until processing. Biological replicates were obtained in triplicate for each paired condition. Proteins were quantified by filter paper assay^86^. 1mg protein per sample was reduced with 10mM TCEP (Tris(2-carboxyethyl) phosphine; Sigma-Aldrich, #646547) at 25 °C for 30 min, alkylated with 55mM CAA (Chloroacetamide, Sigma-Aldrich, #C0267) in darkness at RT for 30 min, and diluted to 1M urea with 100 mM TEAB (Tetraethylammonium bromide; Supelco, #18597) pH 8.2. Proteins were digested with LysC (Wako chemicals, #129-02541) with enzyme:protein ratio of 1:50 (w/w) for 4 h at 37 °C, followed by trypsin (Promega, #V5117) 1:100 (w/w) overnight at 37 °C. Proteins were further digested with trypsin 1:200 (w/w) for 1 h at 37 °C and acidified to a final concentration of 1% TFA (trifluoroacetic acid; VWR Chemicals, #153112E). Acidified peptides were desalted on Affinisep AttractSPE® HLB Disks Spin columns (Affinisep, # Spin15-HLB.T3.50) as follows. Each column was washed with 1 ml 100% ACN (acetonitrile; Biosolve, # 012078) and pre-equilibrated with 2 ml 0.1% TFA before loading the samples. After binding, samples were washed with 500 µl 0.1% TFA and eluted twice with 250 µl 65% ACN, 1% TFA in water. Peptides were snap frozen in liquid nitrogen, lyophilized and keep at −20 °C. Frozen, lyophilized samples were sent to the VIB Proteomics Core (Gent, Belgium) for further processing and LC-MS/MS acquisition. Lyophilized peptides were re-dissolved in water/ACN/TFA (99.4:0.5:0.1, v/v/v) and 1/20 (∼50 µg) of each sample was isolated for shotgun analysis, while 1/2 of the remaining peptides (∼500 µg) were used for phosphopeptide enrichment. Peptides for shotgun analysis were desalted on a reversed phase OMIX C18 tip (Agilent, #A57003100). The tip was first washed 3 times with 100 µl pre-wash buffer (0.1% TFA in water/ ACN (20:80, v/v)) and pre-equilibrated 5 times with 100 µl of wash buffer (0.1% TFA in water) before the sample was loaded on the tip. After peptide binding, the tip was washed 3 times with 100 µl of wash buffer and peptides were eluted twice with 100 µl elution buffer (0.1% TFA in water/ACN (40:60, v/v)). The combined elutions were transferred to HPLC inserts and dried in a vacuum concentrator. Phosphopeptides were enriched with MagReSyn® Ti-IMAC HP beads (ReSyn Biosciences, #MR-THP005) according to the manufacturer’s instructions with slight modifications. Briefly, 100 µl beads (per sample) were washed twice with 70% EtOH, once with 1% NH4OH and three times with a mixture of water/ACN/TFA (14:80:6, v/v/v). Next, the digested sample was incubated with the washed beads for 30 min at RT, the beads were washed once with a mixture of water/ACN/TFA (14:80:6, v/v/v) and three times with a mixture of water/ACN/TFA (19:80:1, v/v/v). Phosphopeptides were eluted from the beads by adding three times 80 µl 1% NH4OH. 60 µl 10% formic acid (FA) was added to the combined eluate and the samples were dried completely in a vacuum concentrator. Dried phosphopeptide samples were dissolved in a mixture of water/ACN/TFA (97.9:2:0.1, v/v/v) and desalted on reversed phase OMIX C18 tips as described for the shotgun samples.

#### LC-MS/MS acquisition

Peptides were re-dissolved in 20 µl (proteome acquisition) or 25 µl (phosphoproteome acquisition) loading solvent A (0.1% trifluoroacetic acid in water/acetonitrile (ACN) (99.5:0.5, v/v)) of which 2 µl (proteome acquisition) or 10 µl (phosphoproteome acquisition) of sample was injected for LC-MS/MS analysis on an Ultimate 3000 ProFlow nanoLC system in-line connected to a MS Orbitrap Exploris 240 mass spectrometer (Thermo) equipped with pneu-Nimbus dual ion source (Phoenix S&T). Trapping was performed at 20 μl/min for 2 min in loading solvent A on a PepMap™ Neo Trap column (Thermo scientific, 300 μm internal diameter (I.D.), 5 μm beads). The peptides were separated on a 250 mm Aurora Ultimate, 1.7µm C18, 75 µm inner diameter (Ionopticks) kept at a constant temperature of 45°C. For proteome acquisition, peptides were eluted by a gradient reaching 26,4 % MS solvent B (0.1% FA in acetonitrile) after 30 min, 44% MS solvent B at 38 min, 56% MS solvent B at 40 min followed by a 5-minutes wash at 56% MS solvent B and re-equilibration with MS solvent A (0.1% FA in water), with a flow rate of 250 nl/min. The mass spectrometer was operated in data-independent (DIA) mode, automatically switching between MS and MS/MS acquisition. Full-scan MS spectra ranging from 375-1500 m/z with a normalized target value of 300%, a maximum fill time of 25 ms and a resolution at of 60,000 were followed by 30 quadrupole isolations with a precursor isolation width of 10 m/z for HCD fragmentation at an NCE of 30% after filling the trap at a normalized target value of 2000% for maximum injection time of 45 ms. MS2 spectra were acquired at a resolution of 15,000 with a scan range of 200-1800 m/z in the Orbitrap analyser without multiplexing. The isolation intervals were set from 400 – 700 m/z with a width of 10 m/z using window placement optimization. For phosphoproteome acquisition, peptides were eluted by a gradient reaching 26,4 % MS solvent B (0.1% FA in acetonitrile) after 80 min, 44% MS solvent B at 95 min, 56% MS solvent B at 100 min followed by a 5-minutes wash at 56% MS solvent B and re-equilibration with MS solvent A (0.1% FA in water), with a flow rate of 250 nl/min. The mass spectrometer was operated in data-dependent (DDA) positive ionization mode, with a cycle time of 1 second. Full-scan MS spectra (350-1200 m/z) were acquired at a resolution of 120,000 in the Orbitrap analyzer after accumulation to a normalized AGC target value of 300% with a maximum injection time of 25 ms. The precursor ions were filtered for charge states (2-6 required), dynamic exclusion (30 s; +/- 10 ppm window, n=1) and intensity (minimal intensity of 8E3). The precursor ions were selected in the quadrupole with an isolation window of 2 m/z and accumulated to a normalized AGC target of 50% or a maximum injection time of 100 ms and activated using HCD fragmentation (30% NCE). The fragments were analysed in the Orbitrap at a resolution of 15,000 and first mass was set to 120. The polydimethylcyclosiloxane background ion at 445.120028 Da was used for internal calibration (lock mass) and QCloud was used to control instrument longitudinal performance during the project.

#### Data analysis

All raw proteome data were processed using FragPipe (version 23.0) with MSFragger (version 4.3) and searched against the reviewed human UniProt/Swiss-Prot proteome (release 2023-01-05, UP000005640), supplemented with reversed-sequence decoys. Searches were performed with with fully tryptic peptides with a maximum of two missed cleavages. Precursor and fragment ion mass tolerances were set to ±10 ppm and 20 ppm, respectively. Carbamidomethylation of cysteine (+57.021 Da) was set as a fixed modification in all searches. Retention time and spectral predictions were generated using MSBooster (version 1.3.9) with DIA-NN-based models to improve PSM rescoring. For DIA proteome quantification, MSFragger was used to generate a spectral library, and label-free quantification (LFQ) was performed using DIA-NN (version 1.8.2), with identifications filtered to 1% FDR at the precursor and protein level. For DDA phosphoproteome quantification, variable modifications included oxidation of methionine (+15.995 Da), N-terminal acetylation (+42.011 Da), and phosphorylation of serine, threonine, and tyrosine residues (+79.966 Da), allowing a maximum of two missed cleavages and three variable modifications per peptide. PSM scoring was performed using Percolator (version 3.7.1) in PSM-only mode with target-decoy competition, and phosphorylation site localization was assessed using PTMProphet (minimum site probability: 0.5; fragment ion tolerance: 10 ppm). Peptide and protein identifications were filtered to 1% and 5% FDR, respectively, using Philosopher (version 5.1.1). Label-free quantification was performed using IonQuant (version 1.11.11) with the MaxLFQ algorithm, inter-sample normalization, and match-between-runs enabled (retention time tolerance: 0.4 min; m/z tolerance: 10 ppm).

Downstream analysis of DIA-LFQ proteome data was performed in R using the FragPipeAnalystR package (version 1.1.0). Quality control was performed by inspection of protein identification numbers, intensity distributions, coefficient of variation, sample-to-sample correlation heatmaps, and PCA. Outlier samples were identified and excluded prior to further analysis. Proteins quantified in at least 66% of samples within at least one experimental condition were retained. The LFQ intensities were log2-transformed, and DEA between palbociclib-treated and vehicle control (DMSO) conditions was assessed using the limma linear model. For gene centric analysis, GSEA was performed using the fgsea package (v. 1.34.2) with MSigDB Hallmark gene sets (minimum gene set size: 10; maximum: 1000), using the log2FC as the ranking metric. Enrichment scores were computed as NES, and significance was assessed by permutation testing. For DDA-LFQ phosphoproteome data, sites with a localization probability below 0.75 were excluded. Sites were retained if detected in at least 50% of replicates within at least one experimental condition. LFQ intensities were log2-transformed, and differential phosphorylation between palbociclib-treated and vehicle control (DMSO) conditions was assessed using limma. Sequence windows of ±7 amino acids centered on the phosphorylated residue were extracted from the UniProt/Swiss-Prot FASTA file used for the database search for downstream analysis. For phosphosite centric analysis, Post-Translational Modification Signature Enrichment Analysis (PTM-SEA) was implemented via the ssGSEA2.0 (version 1.0.0) package with the PTMsigDB v2.0.0 signature database. Sites were ranked by their limma moderated t-statistic. Enrichment scores were computed as NES, and significance was assessed by permutation testing. A minimum overlap of 5 sites per signature was required, and signatures with FDR < 0.1 were considered significant. Kinase activity was predicted with kinase-substrate enrichment analysis (KSEA) using the KSEAapp (version 2.0) package and site-kinase associations derived from the NetworKIN and PhosphoSitePlus databases (July 2016 release). Predicted kinase activity scores were computed as z-score from DEA fold changes and p-values, and z-score-derived p-values were corrected for multiple testing with Benjamini-Hochberg method. A minimum overlap of 5 sites per signature was required, and signatures with FDR < 0.1 were considered significant. All analyses were performed with R (v 4.4.3).

### Signature-based drug search

Signature-based drug search was performed using the Connectivity Map (CMap) database accessed via CLUE from the Broad Institute^88^. The CMap comprises over 1.5 million transcriptional signatures profiling the response of diverse human cancer cell lines to chemical and genetic perturbations. Two gene signatures were queried against the full CMap database: (i) upregulated genes shared across all resistant clones (FC > 1.5, adjusted p-value < 0.1), and (ii) upregulated genes shared across all parental cell lines following palbociclib treatment (FC > 1.5, adjusted p-value < 0.1). Similarity between our signatures and each CMap profile was quantified using the CMap connectivity score (*tau*), a standardized metric ranging from −100 to 100 for lowest to highest similarity, respectively. For signature (i), perturbagens were ranked by most dissimilar connectivity (most negative *tau* scores), to identify compounds capable of transcriptionally reversing the resistance phenotype. For signature (ii), perturbagens were ranked by most similar connectivity (most positive *tau* scores), to identify compounds that recapitulate the transcriptional effects of palbociclib treatment. In both cases, perturbagens were ranked by their *General* score, defined as the median *tau* score across all queried cell lines.

### Synergy assay

Parental and -RP2 cell lines were seeded in 96-well plates at densities ranging from 5 × 10^2^ to 2 × 10^3^ cells per well. After 24 hours, cells plates were treated in a 6 × 5 concentration matrix combining five inhibitors (apicidin, entinostat, GW-843682X, SB-216763, or vorinostat) with either cisplatin or palbociclib. Concentration ranges were selected to span the IC50 of each agent as determined from monotherapy experiments: apicidin and GW-843682X, 6–600 nM; entinostat and vorinostat, 20–2000 nM; SB-216763, 60–6000 nM; cisplatin, 60–5000 nM; and palbociclib, 60–4000 nM. Treatments were maintained for 6 days with a refresh at day 3. Cell viability was assessed by MTT assay as previously described. Drug interactions were quantified using SynergyFinder+^39^, applying the Bliss independence model for cisplatin-based combinations and the Zero Interaction Potency (ZIP) model for palbociclib-based combinations. Synergy scores above 10 were considered synergistic, scores between 0 and 10 additive, and scores below 0 antagonistic. Combination efficacy was calculated with the combination sensitivity score (CSS). All experiments were performed in biological triplicate. Cell viability (%) data of selected combinations are presented as mean ± SD.

### Statistical analysis

All experiments were conducted with a minimum of three independent biological replicates, and data is presented as mean ± SD. Statistical analyses of functional assays were performed using GraphPad Prism version 8.0 and the detailed statistical methods are provided in the respective figure legends and methods section. Prior to statistical analysis, data were assessed for normality using the Shapiro-Wilk test and Q-Q plots. Statistical significance was determined using a threshold of p-value < 0.05. Statistical analyses of multi-omics data were performed using R (v4.4.3). Statistical significance was determined using a threshold of FDR < 0.1.

## Data Availability Statement

All raw RNA-seq data from resistant cell lines generated in this article have been deposited in the Gene Expression Omnibus (GEO) under the accession number [**GSE338897**]. All raw RNA-seq data from parental cell lines had previously been generated by F.M.^8^ and are publicly accessible from the GEO under the accession number GSE312063. All raw proteomic and phosphoproteomic data from parental and resistant cell lines generated in this article have been deposited online on PRIDE server under the accession number [PXD081191 & PXD081216 for the proteome and phosphoproteome, respectively]*. Processed data are provided with this paper.*\*Data will be publicly accessible upon publication*.

## Acknowledgments

This work was supported by Fonds Paul Genicot (recipient: X.B.), Fonds Gaston Ithier (recipient: X.B), Fonds de la Recherche Scientifique - FNRS (CDR n°40008450 and 40028133, recipient: X.B.) and Fondation contre le Cancer (2022–161, recipient: X.B.) B.B. is a Senior Research Associate of the Fonds de la Recherche Scientifique – FNRS. M.A., F.M. and D.J. are supported by Fonds pour la Formation à la Recherche dans l’Industrie et dans l’Agriculture (FRIA, F.R.S.-FNRS) fellowship. M.A. and F.M. are supported by Fonds David et Alice Van Buuren and Fondation Jaumotte-Demoulin. A.V. was recipient of a scholarship from Erasmus+ program. We thank Virginie Imbault for her technical guidance and for ensuring the smooth operation of the laboratory equipment. We thank Anne Lefort, Frédérick Libert, Sabine Paternot and the Brussels Interuniversity Genomics High Throughput BRIGHTcore (www.brightcore.be) for the RNA sequencing. We thank Simon Devos and the VIB Proteomics Core (PRC, www.proteomicscore.sites.vib.be) for the phosphoproteomics enrichment and the HPLC-MS/MS acquisition.

## Author contributions

**M.A.:** Conceptualization, data curation, data analysis, data visualization, writing, reviewing and editing.

**A.V.:** Data curation.

**F.M.:** Data curation, data analysis.

**D.J.:** Data curation.

**B.B.:** Resources, reviewing, and editing.

**X.B.:** Conceptualization, resources, funding acquisition, project administration, writing, reviewing and editing.

All authors read and approved the final manuscript.

Supplementary Figures

---

**Supplementary Figure 1.**
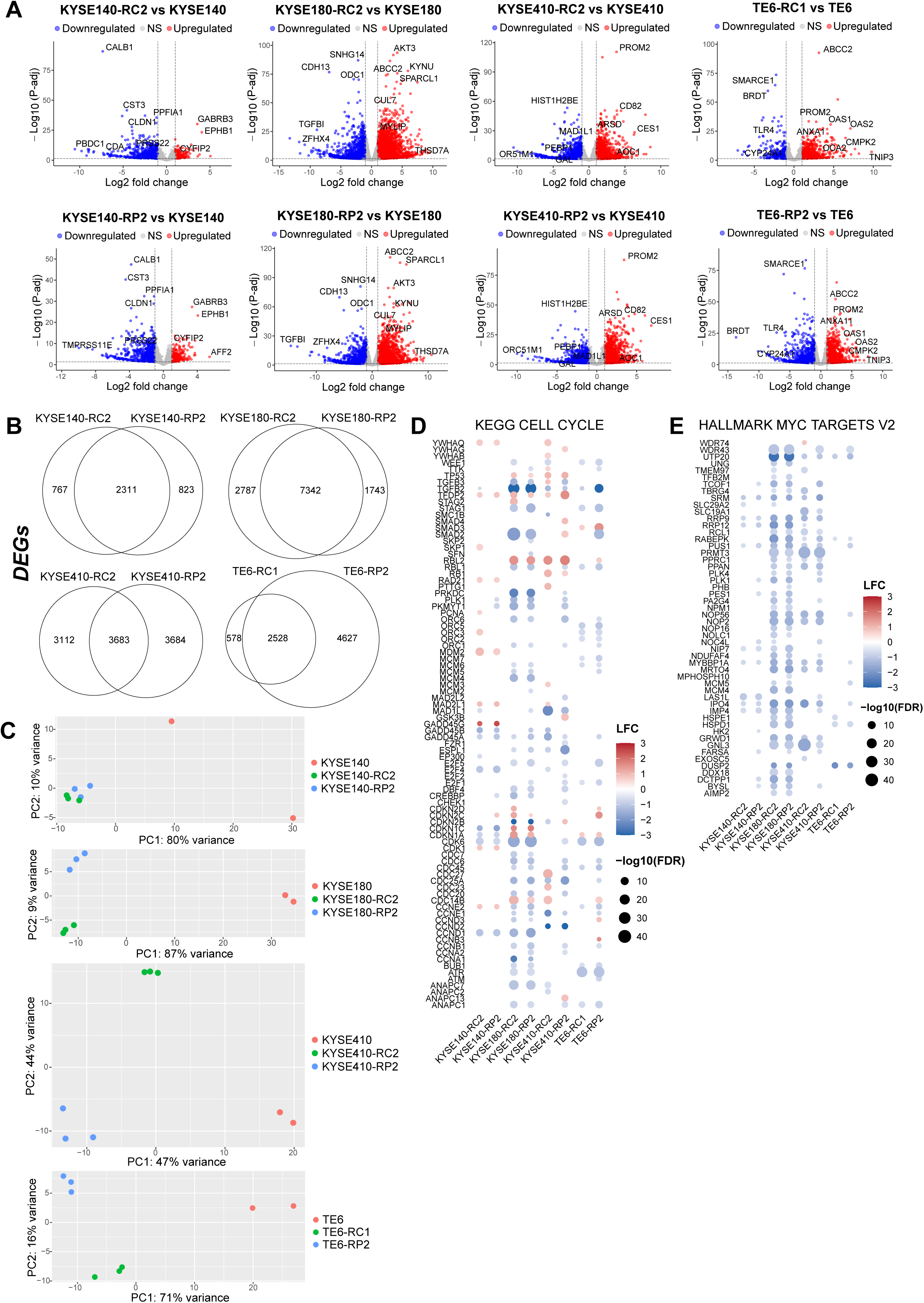
**(A)** Volcano plots of RNAseq data from parental and resistant cell lines. Significance threshold for DEGs was |LFC| > 0.585 and p-adj < 0.1. **(B)** Venn diagrams of RNAseq data from parental and resistant cell lines. Significance threshold for DEGs was |LFC| > 0.585 and p-adj < 0.1. **(C)** Principal component analysis (PCA) of RNAseq data from parental and resistant cell lines (top 500 genes) **(D)** RNA-seq dotplot of resistant versus parental cell lines, showing significant differentially expressed genes related to the cell cycle (Hallmarks gene set). Significance threshold was |LFC| > 0.585 and p-adj < 0.1. **(E)** RNA-seq dotplot of resistant versus parental cell lines, showing significant differentially expressed genes related to MYC targets (Hallmarks gene set). Significance threshold was |LFC| > 0.585 and p-adj < 0.1.

**Supplementary Figure 2.**
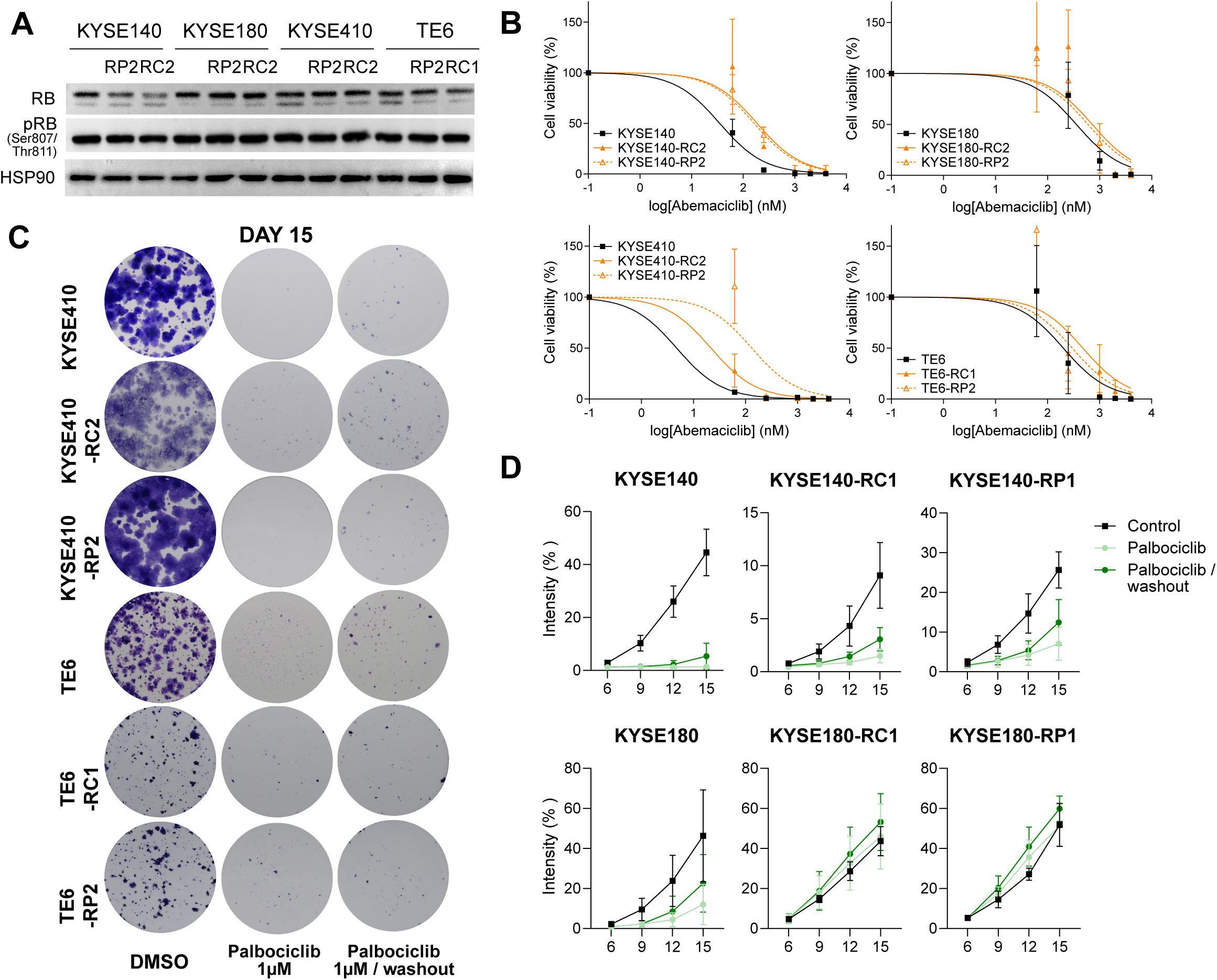
**(A)** Immunoblotting of cell cycle-related proteins RB and its phosphorylated form, p-RB (Ser807/811) in parental and resistant cell lines. Loading control with HS90. **(B)** Viability curves of cell lines treated with abemaciclib (CDK4/6 inhibitor) for 15 days followed by clonogenic assay. Data are mean ± SD. **(C)** Representative images at day 15 of parental and cisplatin-resistant KYSE410 and TE6 cell lines treated with 1μM palbociclib for 15 days, or treated for 6 days with a 9-days washout period (n= 3). **(E)** Difference in growth curves overtime between parental and early resistant (R1) cell lines treated with 1μM palbociclib for 15 days, or treated for 6 days with a 9-days washout period (n= 3). Cell growth is represented by intensity of staining per pixel. Data are mean ± SD (n = 3).

**Supplementary Figure 3.**
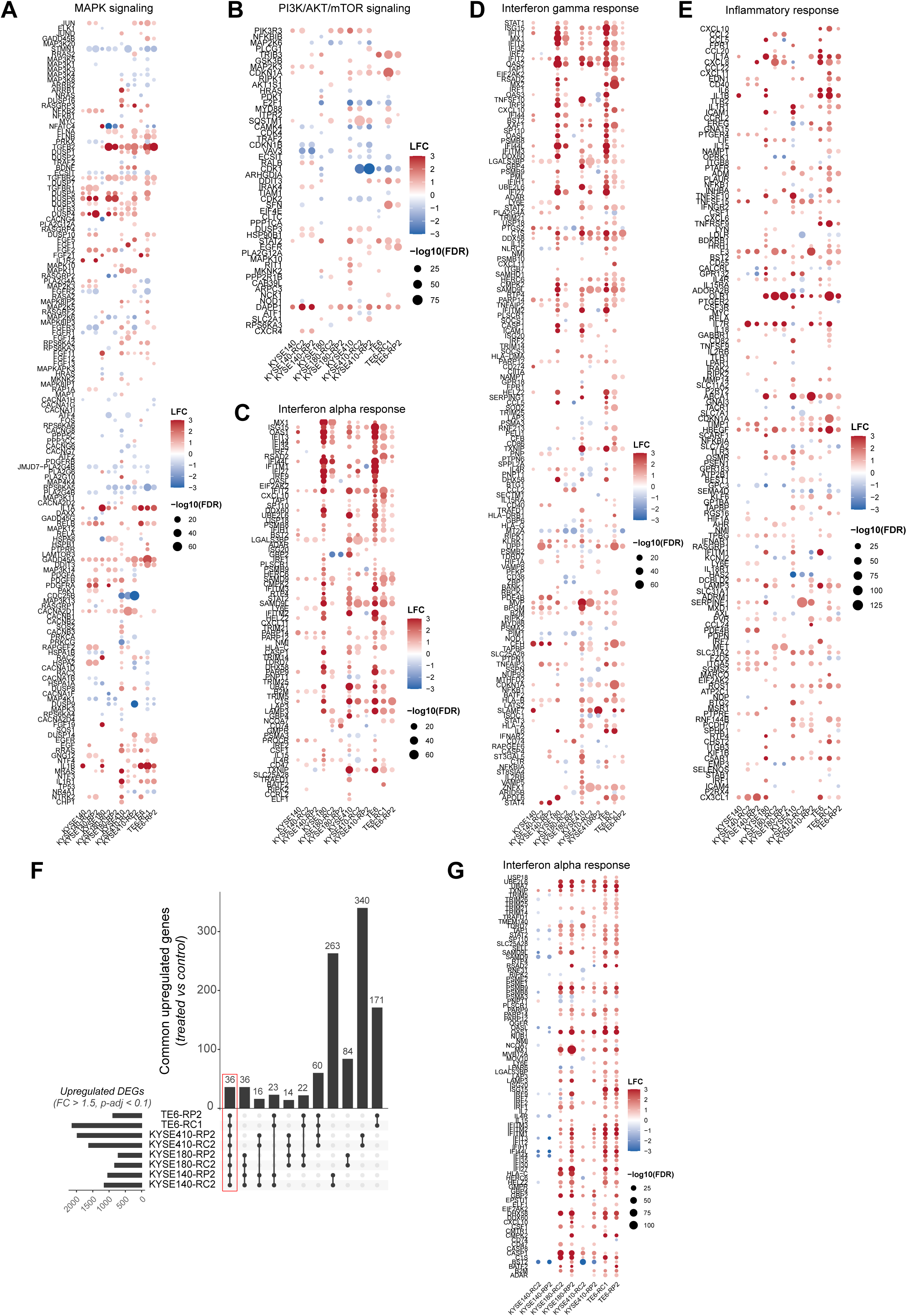
**(A)** RNA-seq dotplot of cell lines treated with palbociclib 1μM for 6 days versus vehicle control (DMSO), showing significant DEGs (|LFC| > 0.585, p-adj < 0.1) related to Hallmark gene set for MAPK signaling and **(B)** PI3K/AKT/mTOR signaling, **(C)** Interferon alpha response, **(D)** Interferon gamma response, **(E)** Inflammatory alpha response. **(F)** Upset plot from RNAseq data of significantly upregulated genes (FC > 1.5, p-adj < 0.1) of cell lines treated with palbociclib 1μM for 6 days versus vehicle control (DMSO). **(G)** RNA-seq dotplot of resistant versus parental cell lines, showing significant DEGs (|LFC| > 0.585 and p-adj < 0.1) related to Interferon alpha response.

**Supplementary Figure 4.**
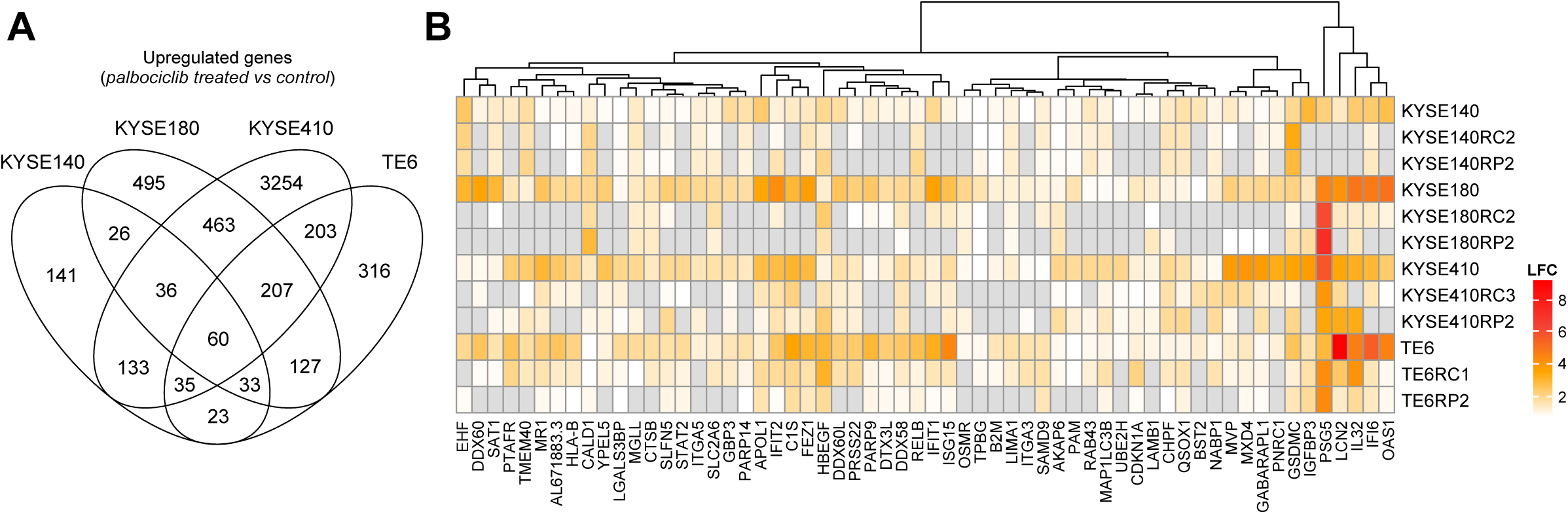
**(A)** Venn diagram from RNAseq data of significant DEGs (FC > 1.5, p-adj < 0.1) of parental lines treated with palbociclib 1μM for 6 days versus vehicle control (DMSO). **(B)** Heatmap of transcriptional signature (i), with DEGs shared by all parental cell lines treated with palbociclib 1μM for 6 days versus vehicle control (DMSO). Significance threshold was defined as FC > 1.5 and p-adj < 0.1.

**Supplementary Figure 5.**
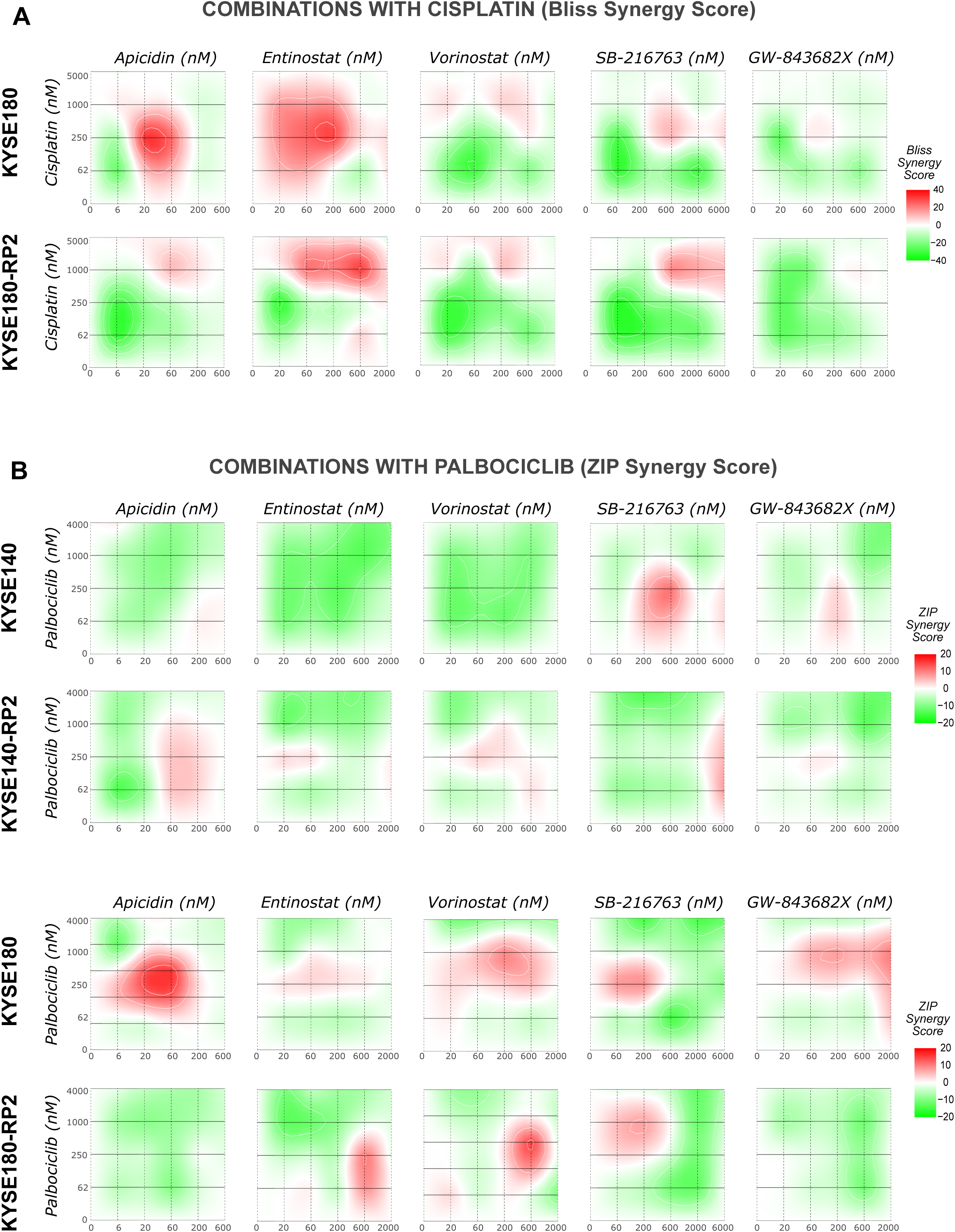
**(A)** Bliss synergy scores heatmaps of KYSE180 and KYSE180-RP2 cell lines following combination of cisplatin with signature-based selected inhibitors. **(B)** ZIP synergy scores heatmaps of KYSE140, KYSE140-RP2, KYSE180 and KYSE180-RP2 cell lines following combination of palbociclib with signature-based selected inhibitors. Synergy scores above 10 were considered synergistic, scores between 0 and 10 additive, and scores below 0 antagonistic.

